# Postsynaptic induction and presynaptic expression of long-term potentiation at excitatory synapses on layer 2/3 VIP interneurons in the somatosensory cortex

**DOI:** 10.1101/2025.07.31.667840

**Authors:** Karolina Bogaj, Joanna Urban-Ciecko

**Author notes:** Corresponding author: Joanna Urban-Ciecko.

## Abstract

Synaptic transmission between specific connection motifs undergoes plastic changes during learning process, however, exact mechanisms underlying synaptic plasticity are still under intense investigation. Long-term potentiation (LTP) of synaptic transmission is a widely used cellular model of synaptic plasticity occurring during learning. Here, we focused on studying LTP at excitatory synapses on layer (L) 2/3 vasoactive intestine polypeptide-expressing interneurons (VIP-INs) in the mouse somatosensory (barrel) cortex. LTP was induced by a pairing protocol of postsynaptic depolarization with extracellular stimulation in acute brain slices of young mice (P21-28). The pairing protocol evoked LTP in L2/3 VIP-INs in control condition, however, pharmacological blocking GABAaR inhibition enhanced LTP. Next, we found that LTP in L2/3 VIP-INs is dependent on metabotropic glutamate receptor type 1 (mGluR-1) and L-type voltage-gated calcium channels (L-type VGCC) but not on NMDARs nor mGluR-5. Here, mGluR-1 acts through G protein-coupled signaling, Src-family pathway, independently of transient receptor potential channels (TRPC). Analyses of paired-pulse ratio (PPR) and coefficient of variation (CV) indicated a presynaptic locus of LTP expression. Presynaptic expression of LTP in VIP-INs relies on retrograde signaling through endocannabinoids (eCBs) but not on brain-derived neurotrophic factor (BDNF). In conclusion, we dissected mechanisms of LTP induction and expression at excitatory inputs to L2/3 VIP-INs in the mouse barrel cortex. LTP at excitatory synapses on VIP-INs might serve as a positive feedback for enhanced VIP-IN-mediated inhibition of SST-INs, leading to disinhibition of excitatory neurons from SST-IN inhibition during learning process.

## 1. Introduction

It has been believed that synaptic plasticity underlies learning process and memory formation (Martin *et al*., 2000). Several lines of evidence indicate that mechanisms underlying synaptic plasticity are cell type-, synapse- and brain region-specific (Maffei *et al*., 2004; Topolnik *et al*., 2006; Banerjee *et al*., 2009; Eng *et al*., 2016; Booker *et al*., 2018; Williams & Holtmaat, 2019; Chistiakova *et al*., 2019; Patton *et al*., 2024). Synaptic plasticity shapes neuronal circuitry, modulating both excitatory (glutamatergic) neurons and inhibitory (GABAergic) interneurons (Wang & Maffei, 2014; Williams & Holtmaat, 2019; Field *et al*., 2020). Neocortical GABAergic interneurons, even though they represent merely 20 % of the entire neuronal population, are immensely divergent group of cells. Vasoactive intestinal polypeptide-expressing interneurons (VIP-INs) constitute around 13 % of all neocortical interneurons (Prönneke *et al*., 2015), but are highly abundant in L2/3 of the barrel cortex (Jiang *et al*., 2015; Prönneke *et al*., 2015). Most of L2/3 VIP-INs are characterized by bipolar morphology with dendritic arborizations in L1 and also across all deeper cortical layers (Jiang *et al*., 2015, 2023; Prönneke *et al*., 2015; Georgiou *et al*., 2022). For this reason, it has been suggested that L2/3 VIP-INs might receive excitatory inputs from local excitatory neurons in all cortical layers as well as from long-range projecting neurons (Lee *et al*., 2013; Prönneke *et al*., 2015; Audette *et al*., 2018; Williams & Holtmaat, 2019; Naskar *et al*., 2021; Georgiou *et al*., 2022; Jiang *et al*., 2023). L2/3 VIP-INs provide disinhibition of glutamatergic neurons from other interneurons (Chéreau *et al*., 2022), especially from somatostatin interneurons (SST-INs) (Lee *et al*., 2013; Jiang *et al*., 2015; Walker *et al*., 2016). VIP-INs are one of the least studied class of interneurons in terms of their plasticity, only a few research groups have reported plasticity in VIP-INs (Canto-Bustos *et al*., 2022; Kanigowski & Urban-Ciecko, 2024, 2025; McFarlan *et al*., 2024b, 2024a), albeit disinhibition mediated by VIP-INs is crucial for plasticity during learning process (Lee *et al*., 2013; Fu *et al*., 2015; Williams & Holtmaat, 2019; Canto-Bustos *et al*., 2022).

Long-term potentiation (LTP) of synaptic transmission is a widely used cellular model of synaptic plasticity underlying learning and memory process (Nicoll, 2017). LTP is a long-lasting strengthening of synapses with various underlying mechanisms of induction and expression (Citri & Malenka, 2008). LTP induction has been observed to be predominantly NMDAR-dependent (Jia *et al*., 1998; Citri & Malenka, 2008; Banerjee *et al*., 2009; Hasan *et al*., 2013; Cui *et al*., 2016; Williams & Holtmaat, 2019; Gómez *et al*., 2021; Hirai *et al*., 2022; Patton *et al*., 2024). Additionally there have been other factors reported that can influence LTP, such as metabotropic glutamate receptors (mGluRs), calcium-permeable AMPARs (CP-AMPARs), or cholinergic receptors (Topolnik *et al*., 2005, 2006; Alkadhi, 2021; McFarlan *et al*., 2023). mGluRs belong to the G protein-coupled receptor family. Class I consists of mGluR-1 and −5 subtypes that may trigger different signaling pathways leading to LTP or long-term depression (LTD) (Perez *et al*., 2001; Benquet *et al*., 2002; Gubellini *et al*., 2003; Topolnik *et al*., 2005, 2006; Nahir *et al*., 2010; Kubota *et al*., 2014; Eng *et al*., 2016). Expression of LTP depends on pre- or postsynaptic mechanisms (Nicoll, 2003; Citri & Malenka, 2008; Yang & Calakos, 2013). Postsynaptic expression involves potentiation of AMPAR activity through receptor phosphorylation or insertion of new receptors to the postsynaptic membrane (Citri & Malenka, 2008; De León-López *et al*., 2025), whereas presynaptic expression of LTP relies on an increase of neurotransmitter release probability (Bender *et al*., 2009; Yang & Calakos, 2013). It has been found that in the motor cortex, LTP at excitatory synapses on L2/3 VIP-INs is expressed postsynaptically but induction mechanisms has not been established (McFarlan *et al*., 2024b).

Here, we described LTP at excitatory synapses on L2/3 VIP-INs in the mouse somatosensory (barrel) cortex. We identified receptors and channels crucial for LTP induction and propose a retrograde signaling factor necessary for LTP expression at these synapses. We found that in the somatosensory cortex LTP induction depends on mGluR-1 and long-lasting voltage-gated calcium channel (L-type VGCC) but not on NMDARs, whereas surprisingly endocannabinoids (eCBs) act as retrograde messengers for presynaptic expression of LTP through activation of the cannabinoid receptor 1 (CBR-1).

## 2. Material and Methods

### 2.1. Ethics statement

All experiments were performed in compliance with the European Community Council Directive (86/609/EEC) and with the Act on the Protection of Animals Used for Scientific or Educational Purposes in Poland (Act of 15 January 2015, changed 17 November 2021; directive 2010/63/EU) considering transgenic mice employment in research. Animal handling and procedures were conducted in accordance with the Institutional Animal Care and Use at the Nencki Institute guidelines.

### 2.2. Animals

Mice strains were purchased from the Jackson Laboratory. Heterozygous VIP-Cre::Ai14 offspring of homozygous VIP-Cre females bred with homozygous Ai14 males (stock #010908 and #007908 respectively) were used (both sexes, at the age of P21-P28). Mice were weaned at P21 and group housed (according to sex) with unlimited food and water availability, in controlled condition of 12-hour day to night cycle, stable temperature and humidity.

### 2.3. Acute brain slices preparation

For brain extraction, animals were humanely killed by decapitation following deep anesthesia with isoflurane. Slices were then obtained in a paracoronal plane to preserve the barrel field architecture (Urban-Ciecko *et al*., 2018; Kanigowski *et al*., 2023; Bogaj *et al*., 2023). Slices were sectioned to 350 μm with Leica Vibratome, in cold artificial cerebrospinal fluid (ACSF). For recovery and recording, slices were maintained in warm ACSF, heated up to 31 °C. The solution contained (mM): 119 NaCl, 2.5 KCl, 1.3 MgSO_4_, 2.5 CaCl_2_, 1 NaH_2_PO_4_, 26.2 NaHCO_3_, 11 glucose and was bubbled with 95/5 % O_2_/CO_2_ (Urban-Ciecko *et al*., 2018; Bogaj & Urban-Ciecko, 2025).

### 2.4. Electrophysiology

Neurons were visualized under water-immersion lens with 40x magnification (Axio Examiner A1, Zeiss). In vitro whole-cell patch-clamp technique was performed from fluorescently-labeled VIP-INs in L2/3 of the mouse somatosensory (barrel) cortex. Recordings were acquired with borosilicate glass electrodes (4-7 MΩ), filled with internal solution composed of (mM): 130 cesium gluconate, 10 HEPES, 0.5 EGTA, 8 NaCl, 10 TEA-Cl, 4 Mg-ATP, and 0.3 Na-GTP, adjusted to pH 7.3 and osmolarity 290 mOsm (Kanigowski *et al*., 2023).

To stimulate excitatory inputs to VIP-INs, bipolar electrode made from theta glass filled with ACSF, was placed in L2/3 below recorded cell (Fig. 1A). Membrane potential of a neuron was held at −65 mV in voltage-clamp mode to acquire evoked excitatory postsynaptic currents (EPSCs). EPSCs were recorded for a stable 5-10 min baseline at the frequency of 0.1 Hz. The intensity of stimulation (Isolator ISO-200, CircleLabs) was set individually for each neuron, to produce minimal stable response to the stimulus lasting 0.2 ms. The EPSC amplitude was analyzed within the area of 3-5 ms after the stimulus artifact. LTP was induced by postsynaptic depolarization to 0 mV paired with the extracellular stimulus, 55-60 consecutive times at the same frequency as for baseline stimulation (Feldman, 2000). Evoked EPSCs were then recorded as in baseline protocol at −65 mV for at least 30 min after the pairing protocol. LTP was assessed 25-30 min after the end of the LTP pairing. For paired-pulse stimulation, 2 pulses were delivered in 50 ms intervals with the frequency of 0.1 Hz (5-10 repetitions after baseline and 30 min after the LTP protocol).

**Fig. 1.**
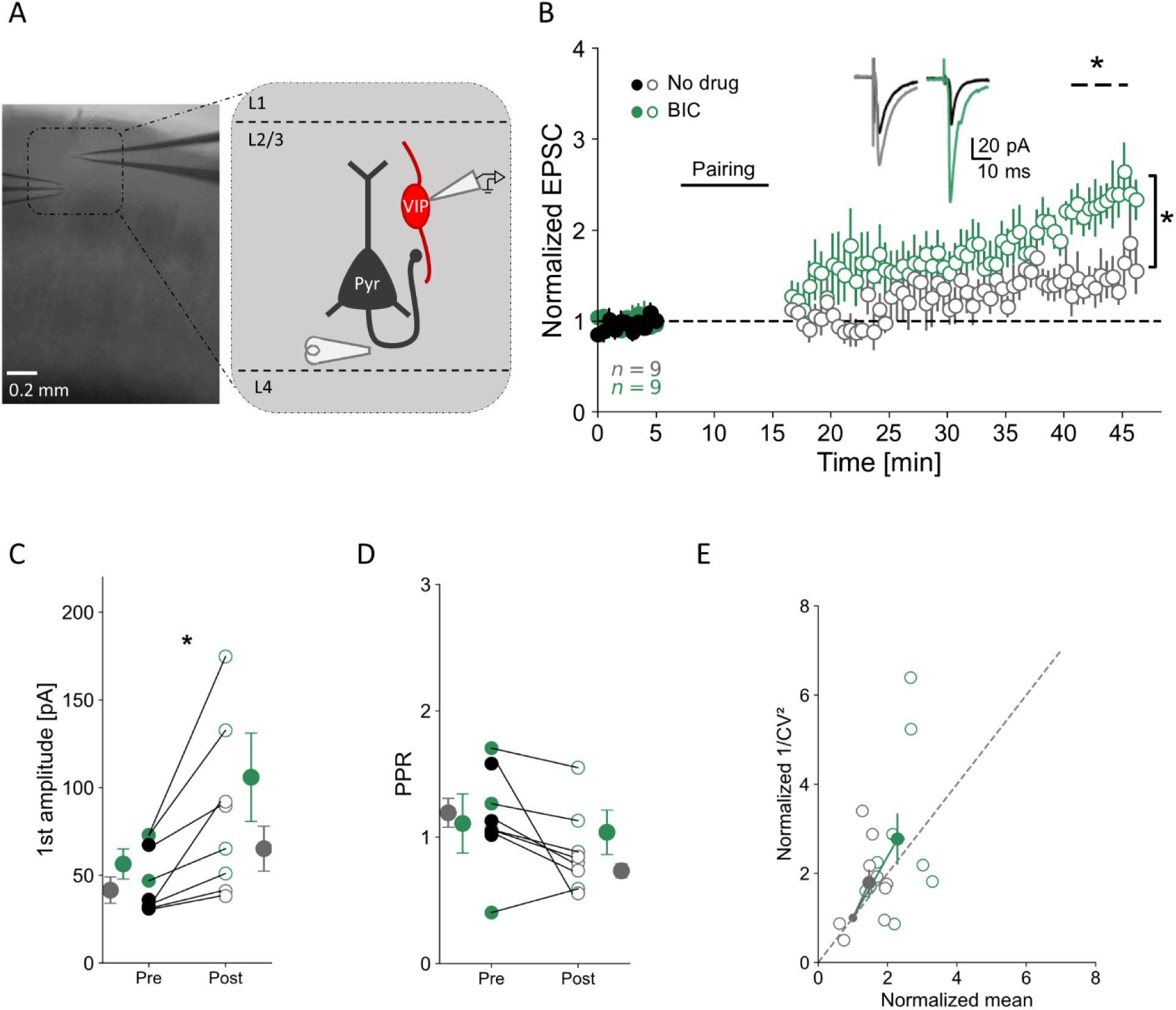
Inhibition of GABAaRs elevates LTP at excitatory inputs to L2/3 VIP-INs in the somatosensory cortex. **A)** Picture of acute brain slice with stimulating and recording electrodes in L2/3 of the mouse barrel cortex, and a schematic illustrating the stimulation paradigm – excitatory inputs from pyramidal cells (Pyr – dark gray) are stimulated to evoke EPSCs in fluorescently-labeled VIP-INs (VIP – red). **B)** Time course plot of normalized EPSC before and after the pairing protocol in control condition (No drug: pre pairing – black circles, post pairing – light gray circles) and in bath application of the GABAaR antagonist (bicuculline, BIC: pre pairing – solid green circles and post pairing – open green circles). Horizontal dashed line with an asterisk indicates statistically significant differences for LTP induction in each experimental condition, vertical line with an asterisk indicates a significant difference between the two recording conditions. Inset shows examples of averaged EPSC traces for pre and post LTP induction protocol. **C)** Averaged 1^st^ amplitude for pre and post LTP induction (pooled data for No drug and GABAaR blockage conditions). **D)** Paired pulse ratio for pre and post LTP induction protocol (pooled data as for C). **E)** Normalized 1/CV^2^ plotted against normalized amplitude for both conditions. Mean ± SEM, * p < 0.05.

Access resistance (R_a_) was monitored throughout recordings. Neurons with a change in R_a_ over 30 % or exhibiting unstable baseline were excluded from further analysis.

Data was acquired using Multiclamp 700B amplifier and digitized with Digidata 1550B (Molecular Devices). Data was filtered at 3 kHz, sampled at 20 kHz and collected by pClamp 10 (Molecular Devices).

### 2.5. In vitro pharmacology

The following drugs were bath applied throughout the recordings: GABAaR antagonist – Bicuculline methiodide, BIC (Sigma-Aldrich), 5 µM; NMDAR antagonist – APV, 50 µM; mGluR-5 antagonist – MTEP, 10 µM; mGluR-1 antagonist – LY367385, 50 µM; inhibitor of Src-family tyrosine kinases – PP2, 25 µM; TRPC antagonist – SKF96365, 10 µM; L-type VGCC antagonist – Nifedipine (Sigma-Aldrich), 10 µM; TrkB receptor antagonist – ANA-12, 10 µM; CBR-1 antagonist – AM-251 (HelloBio), 3 or 10 µM; A2R antagonist – SCH58261, 100 nM; and A1R antagonist – DPCPX, 100 nM. Slices were washed with drug solution for at least 10 min prior to data acquisition. GDPβS (Jena Bioscience), 1mM was applied in internal solution replacing GTP. Lipase inhibitor – THL (HelloBio), was dissolved in DMSO and further diluted in internal solution (10 µM final concentration). DMSO final concentration was 1µl /1ml. In the experiments with pharmacological agents added to internal solution, the baseline collection started at least 10 minutes after cell break-in to allow for the effective perfusion of internal solution to the cell.

Unless indicated otherwise, all drugs were acquired from Tocris Bioscience.

### 2.6. Data analysis

Dataset was analyzed with Clampfit software (Molecular Devices) and in Python environment. The EPSC amplitude was calculated within 3-5 ms after stimulation artifact and normalized to the mean baseline events. Paired pulse ratio (PPR) was assessed as EPSC_2_/EPSC_1_ for baseline and 30 min post-LTP protocol. Coefficient of variation (CV) was analyzed as standard deviation of amplitude divided by averaged amplitude of 30 consecutive sweeps. Then inverted squared CV (1/CV^2^) was calculated, and normalized 1/CV^2^ against normalized mean amplitude was plotted to assess pre- or postsynaptic locus of LTP expression (Brock *et al*., 2020). If averaged dataset lays on the unity line (y = x), expression of LTP occurred due to mixed pre- and postsynaptic changes, whereas mean datapoint below or above y = x line indicates postsynaptic or presynaptic locus of plasticity expression, respectively.

### 2.7. Statistical analysis

Data is presented as mean ± standard error of mean (SEM). Normal distribution was checked using Shapiro-Wilk test. For analysis of two groups, paired t-test, Wilcoxon test or independent t-test were used, as noted. To evaluate LTP induction in individual recordings, paired t-test was used on raw data and success rate of LTP was calculated as % of recordings in which an increase in the amplitude was statistically significant. To assess LTP induction in a given condition, one sample t-test was used on normalized data. Difference of p < 0.05 was considered statistically significant. Statistics and data visualization were obtained with Python.

## 3. Results

### 3.1. Pharmacological blockage of GABAaRs facilitates LTP at excitatory synapses on L2/3 VIP-INs

To study synaptic plasticity of excitatory synapses on L2/3 VIP-INs in the somatosensory cortex, we performed whole-cell patch-clamp recordings in acute brain slices (Fig. 1A) obtained from transgenic mice with VIP-INs expressing td-Tomato fluorescent protein. LTP was evoked by extracellular electrical stimulation of excitatory terminals paired with membrane depolarization of a patched cell (Feldman, 2000). Stimulating electrode was placed in L2/3 below recorded interneurons (Fig. 1A) to recruit local inputs (Porter *et al*., 1998).

First, we performed the pairing protocol in control condition with no drugs administered. We managed to trigger LTP in 9 out of 9 cells (100 % success rate); the EPSC amplitude increased by 46 % when measured 30 min after the induction protocol (Fig.1B, normalized amplitude: 1.46 ± 0.17, p = 0.034, n = 9, one sample t-test). Since blocking GABAaR-mediated inhibition facilitates LTP induction in many brain regions (Grover & Yan, 1999), we then asked whether inhibition mediated via these receptors influences LTP induction at excitatory synapses on L2/3 VIP-INs. Bath application of the GABAaR antagonist (BIC) evoked LTP in 9 out of 9 cells (100 % LTP success rate, Fig. 1B, normalized amplitude: 2.32 ± 0.21, p < 0.001, n = 9, one sample t-test) and, indeed, significantly enhanced LTP in comparison to no drug condition (p = 0.008, independent t-test). For this reason, all the following experiments were performed in the presence of BIC in ACSF.

To characterize a locus of LTP expression, we analyzed PPR and 1/CV^2^ (McFarlan *et al*., 2024b). We observed a significant increase in 1^st^ amplitude (Fig. 1C, pooled data, pre 48.99 ± 6.72 pA vs. post 85.55 ± 16.94 pA, p = 0.017, n = 8, paired t-test), and a reduction in PPR after LTP induction (Fig. 1D, pooled data, pre 1.15 ± 0.14 vs. post 0.89 ± 0.114, p = 0.055, n = 8, Wilcoxon test), together suggesting an increase of glutamate release probability. Furthermore, 1/CV^2^ analysis showed possible presynaptic locus of LTP expression, since most of the datapoints are scattered above the unity line indicating alterations in the presynaptic site (Fig. 1E).

### 3.2. LTP at excitatory synapses on VIP-INs is NMDAR independent

Next, we aimed to uncover mechanisms of LTP induction. Since the protocol evoking LTP relies on membrane depolarization, we searched for glutamate- and voltage-dependent mechanisms. We rejected the possibility that CP-AMPA receptors mediate LTP, because depolarization to 0 mV leads to channel pore blockage by endogenous polyamines (Bowie & Mayer, 1995; Topolnik *et al*., 2005). Also, it has been shown that CP-AMPARs do not participate in LTP in VIP-INs in the motor cortex (McFarlan *et al*., 2024a). However, it has been well established that NMDARs are voltage-sensitive (Nowak *et al*., 1984; Mayer *et al*., 1984; Vargas-Caballero & Robinson, 2004), hence we blocked those receptors with APV to assess their putative role in induction of LTP at excitatory synapses on L2/3 VIP-INs. Surprisingly, the NMDAR antagonist failed to prevent LTP induction at these synapses in 10 out of 10 cases (100 % LTP success rate, Fig. 2A, normalized amplitude 1.85 ± 0.17, p = 0.001, n = 10, one sample t-test). Moreover, further analysis showed enhancement of the first amplitude (Fig. 2B, pre 66.35 ± 2.67 pA vs. post 121.24 ± 17.59 pA, p = 0.025, n = 6, paired t-test), diminished PPR (Fig. 2C, pre 1.28 ± 0.08 vs. post 1.09 ± 0.05, p = 0.049, n = 6, paired t-test) and increased 1/CV^2^ (Fig. 2D), hinting towards a presynaptic locus of LTP expression.

**Fig. 2.**
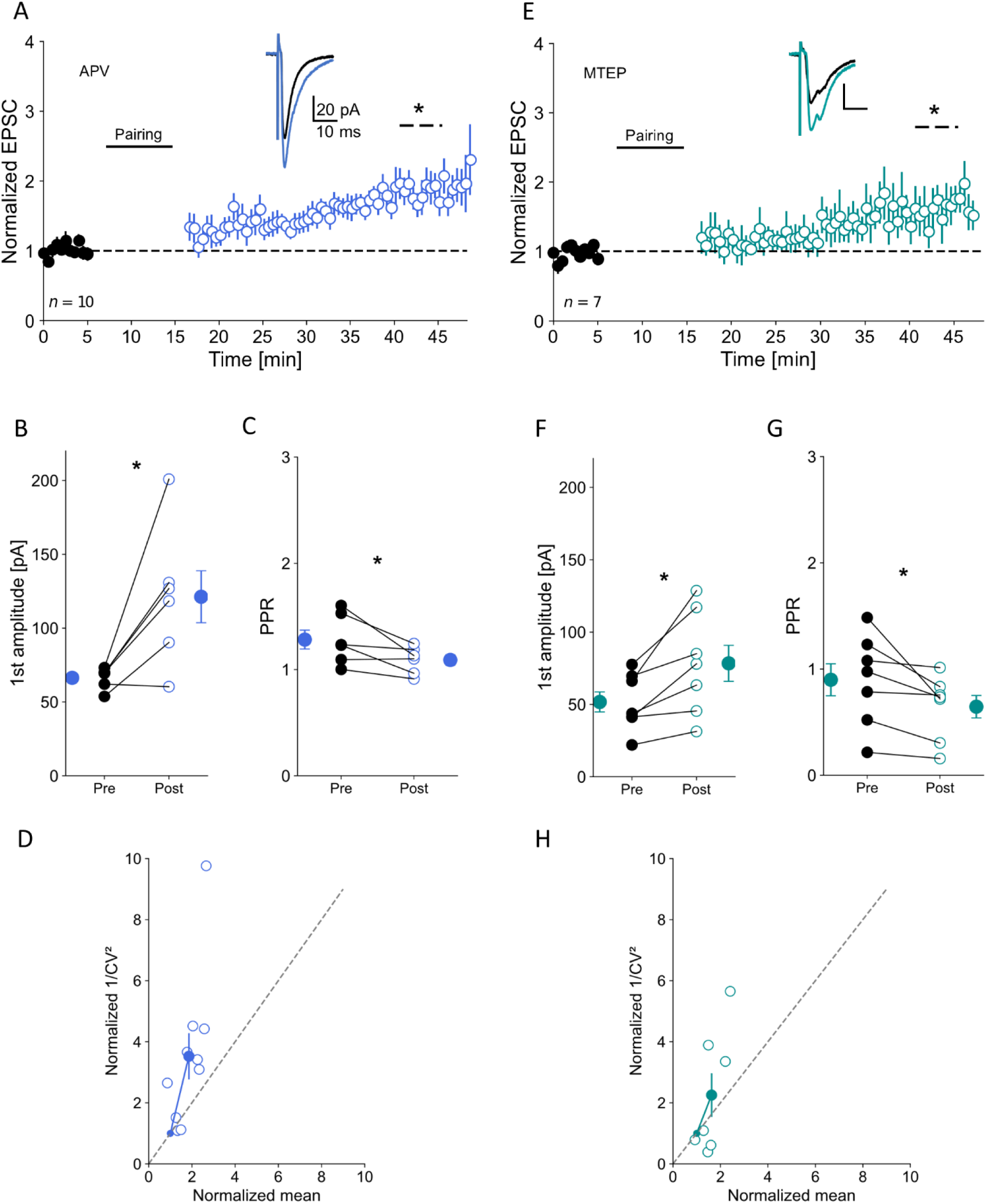
LTP on L2/3 VIP-INs is NMDAR and mGluR-5 independent. **A)** Time course plot of normalized EPSC before and after the pairing protocol in the presence of the NMDAR antagonist (APV); pre pairing – black circles, post pairing – blue circles. Inset: example EPSC traces for pre and post pairing. **B)** Averaged 1^st^ amplitude for pre and post LTP induction protocol with bath applied NMDAR antagonist. **C)** Averaged paired pulse ratio for pre and post LTP induction in APV. **D)** 1/CV^2^ analysis in APV condition. **E)** Same as in A) but for mGluR-5 blockage (MTEP) condition (pre pairing – black circles, post pairing – turquoise circles). **F-H)** Same as B-D) but with bath applied MTEP. Mean ± SEM, * p < 0.05.

### 3.3. Postsynaptic mGluR-1 is responsible for LTP in VIP-INs

After establishing that NMDARs are not involved in LTP induction at excitatory synapses on L2/3 VIP-INs, we searched for other neurotransmitter targets. Metabotropic glutamate receptors (mGluR-1 and mGluR-5) are expressed in cortical neurons (Kerner *et al*., 1997; Heidinger *et al*., 2002) and might be responsible for LTP induction (Huemmeke *et al*., 2002; Topolnik *et al*., 2006). Blocking mGluR-5 with MTEP did not abolish LTP after the pairing protocol in 6 out of 7 cells (86% LTP success rate, Fig. 2E, normalized amplitude 1.64 ± 0.21, p = 0.031, n = 7, one sample t-test; Fig. 2F, amplitude pre 51.7 ± 6.94 pA vs. post 78.41 ± 12.44 pA, p = 0.014, n = 7, paired t-test). Changes in PPR (Fig. 2G, pre 0.89 ± 0.15 vs. post 0.64 ± 0.11, p = 0.042, n = 7, paired t-test) and 1/CV^2^ (Fig. 2H) still suggested the presynaptic locus of LTP expression. However, the pairing protocol failed to induce LTP in VIP-INs in the presence of the mGluR-1 antagonist (LY367385). In this case, LTP was induced in 3 out of 9 cells (33 % LTP success rate, Fig. 3A, normalized amplitude1.28 ± 0.25, p = 0.313, n = 9, one sample t-test; Fig. 3B, amplitude pre 64.6 ± 8.27 pA vs. post 62.13 ± 9.33 pA, p = 0.700, n = 9, paired t-test). Also, there were no changes in PPR (Fig. 3C, pre 1.10 ± 0.11 vs. post 1.36 ± 0.17, p = 0.185, n = 9, paired t-test) and 1/CV^2^ (Fig. 3D) in this condition.

**Fig. 3.**
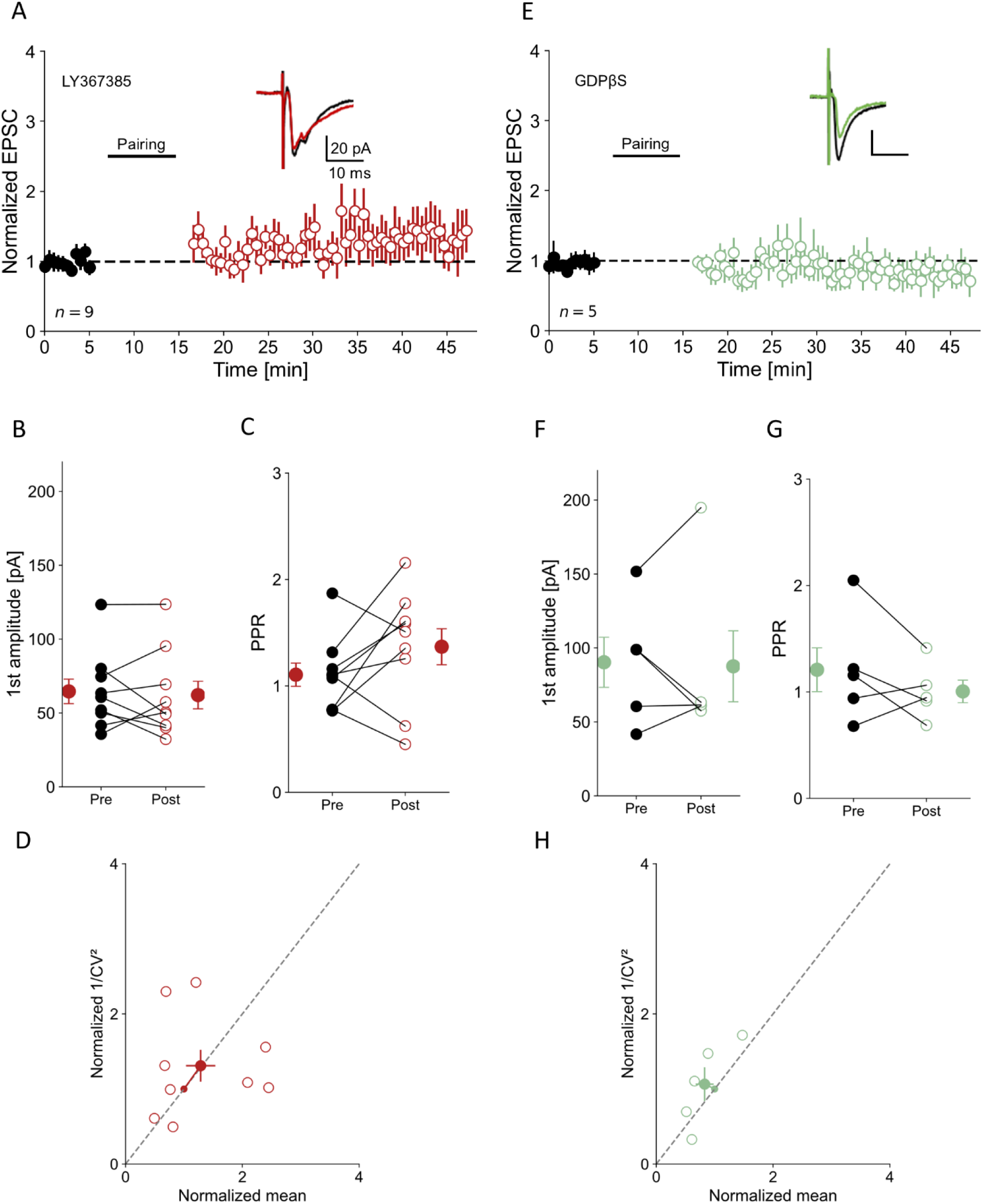
Induction of LTP at excitatory synapses on L2/3 VIP-INs is mediated by postsynaptic mGluR-1 through G protein-coupled signaling pathway. **A)** Normalized EPSC in bath-applied mGluR-1 antagonist (LY367385) condition (pre pairing – black circles, post pairing – red circles). Inset: examples of averaged EPSC traces for pre and post pairing. **B)** Averaged 1^st^ amplitude for pre and post LTP induction protocol in the presence of LY367385. **C)** Averaged paired pulse ratio pre and post LTP induction with impeded mGluR-1. **D)** 1/CV^2^ analysis in LY367385. **E)** Same as A) but with GDPβS in internal solution (pre pairing – black circles, post pairing – light green circles). **F-H)** Same as B-D) but in GDPβS. Mean ± SEM.

Since mGluRs are G protein-coupled receptors but can be expressed both pre- and postsynaptically (Falcón-Moya *et al*., 2020) and astrocytes also can be involved in signaling through mGluRs (Kofuji & Araque, 2021), we asked whether mGluRs-1 modulating LTP in VIP-INs are located postsynaptically. To address this question, we loaded VIP-INs with GDPβS, an inhibitor of G protein-coupled signaling pathways. GDPβS blocked LTP induction in 4 out of 5 cells (20 % LTP success rate) indicating the postsynaptic action of mGluRs-1 in this process (Fig. 3E, normalized amplitude 0.83 ± 0.15, p = 0.374, n = 5, one sample t-test; Fig. 3F amplitude pre 90.30 ± 16.93 pA vs. post 87.59 ± 23.99 pA, p = 0.874, n = 5, paired t-test; Fig. 3G PPR pre 1.21 ± 0.20 vs. post 1.01 ± 0.11, p = 0.305, n = 5, paired t-test; Fig. 3H 1/CV^2^).

mGluR-1 can act through two independent mechanisms in order to produce calcium influx: through activation of Src-family kinases or protein kinase C (PKC) (Topolnik *et al*., 2006; Eng *et al*., 2016; Sugawara *et al*., 2017). To reveal the mode of mGluR-1 action, we applied Src tyrosine kinase antagonist (PP2). In this condition, we could not elicit LTP in VIP-INs (LTP was induced in 2 out of 7 cells, 29 % LTP success rate, Fig. 4A, normalized amplitude 1.04 ± 0.17, p = 0.255, n = 7, one sample t-test; Fig. 4B, amplitude pre 80.01 ± 3.20 pA vs. post 87.41 ± 19.50 pA, p = 0.688, n = 7, Wilcoxon test). There were also no changes observed in PPR analysis (Fig. 4C, pre 1.16 ± 0.09 vs. post 1.10 ± 0.05, p = 0.669, n = 7, paired t-test), nor in 1/CV^2^ (Fig. 4D).

**Fig. 4.**
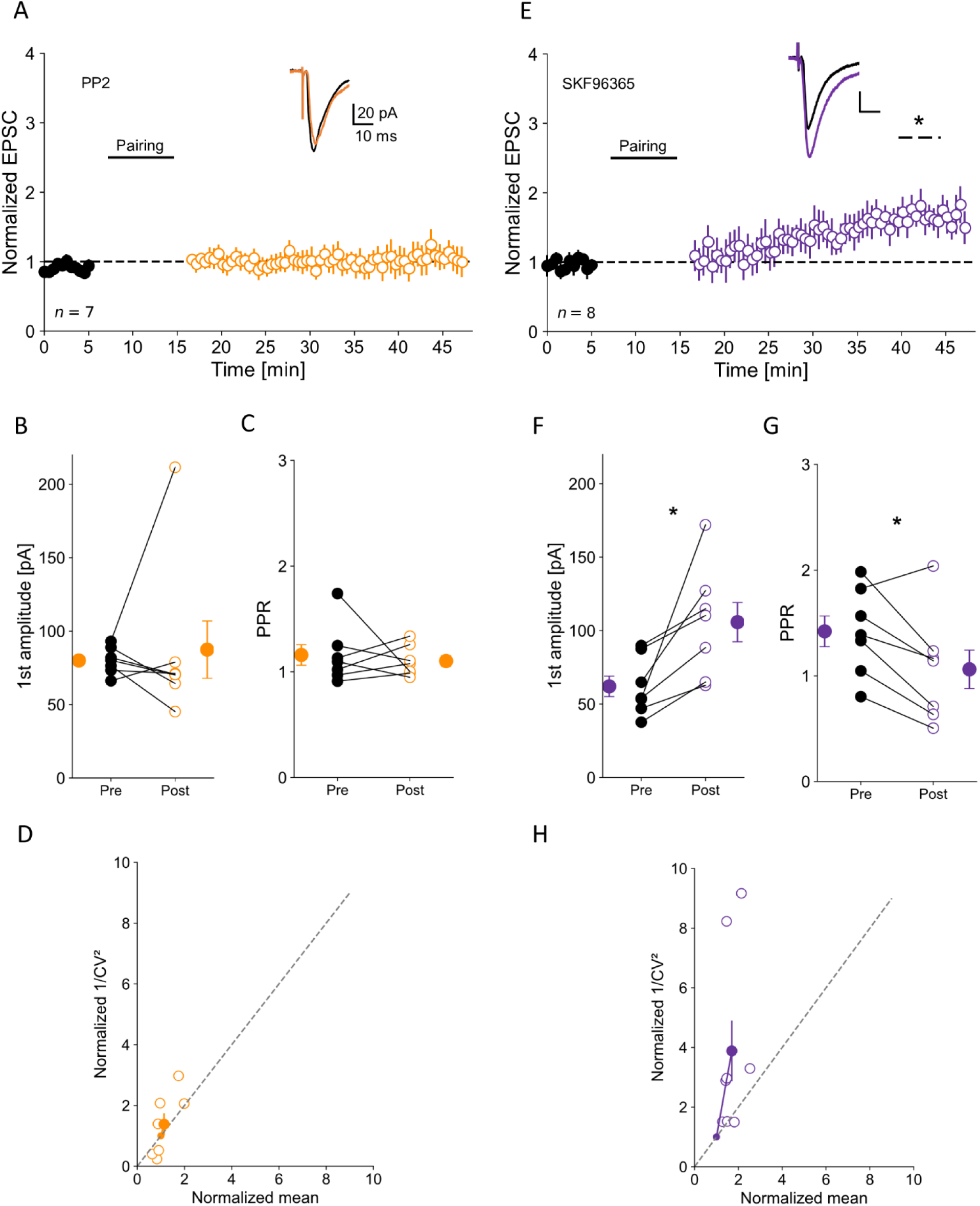
Src-kinase is necessary for LTP in VIP-INs but works through a TRPC-independent mechanism. **A)** Inhibition of Src-kinase abolishes LTP. Normalized EPSC after bath application of Src-kinase inhibitor (PP2); pre pairing – black circles, post pairing – orange circles. Inset: EPSC examples for pre and post LTP induction. **B)** Averaged 1^st^ amplitude for pre and post pairing protocol with inhibited Src-kinase. **C)** Averaged paired pulse ratio for pre and post pairing in PP2 administered condition. **D)** 1/CV^2^ analysis in PP2. **E)** TRPCs are not involved in LTP as shown by time course plot of normalized EPSC before and after the pairing protocol in the presence of TRPC blocker (SKF96365, pre pairing – black circles, post pairing – purple circles). **F-H)** Same as B-D) but in SKF96365. Mean ± SEM, * p < 0.05.

Further, we wanted to know whether the Src pathway leads to activation of transient receptor potential channels (TRPCs) (Topolnik *et al*., 2006; Kubota *et al*., 2014). Blocking TRPCs with SKF96365 did not affect LTP induction in 8 out of 8 cells (100 % LTP success rate, Fig. 4E, normalized amplitude 1.65 ± 0.17, p = 0.008, n = 8, one sample t-test). The EPSC amplitude increased nearly 2-fold post pairing protocol (Fig. 4F, pre 62.06 ± 6.98 pA vs. post 105.78 ± 13.37 pA, p = 0.016, n = 7, Wilcoxon test). PPR significantly decreased (Fig. 4G, pre 1.42 ± 0.14 vs. post 1.06 ± 0.18, p = 0.022, n = 7, paired t-test) and 1/CV^2^ analysis (Fig. 4H) lies above the unity line, showing possible presynaptic locus of synaptic changes.

As we found that TRPCs do not mediate LTP induction, we aimed to find other potential targets for Src kinase. We targeted L-type voltage-gated calcium channels (L-type VGCC) as they can be coupled with mGluRs (Chavis *et al*., 1996, 1998; Topolnik *et al*., 2009; Booker *et al*., 2018) and activity of these channels can be potentiated by tyrosine kinase family (Man *et al*., 2023). Moreover, the pairing protocol involves membrane depolarization, which presumably activates VGCC. Indeed, the L-type VGCC blocker (nifedipine) abolished LTP induction in 5 out of 5 VIP-INs (0 % LTP success rate, Fig. 5A, normalized amplitude 0.89± 0.06, p = 0.192, n = 5, one sample t-test; Fig. 5B, amplitude pre 107. 58 ± 28.40 pA vs. post 93.54 ± 22.48 pA, p = 0.147, n = 4, paired t-test; Fig. 5C, PPR pre 0.81 ± 0.13 vs. post 0.83 ± 0.04, p = 0.870, n = 4, paired t-test; Fig. 5D 1/CV^2^).

**Fig. 5.**
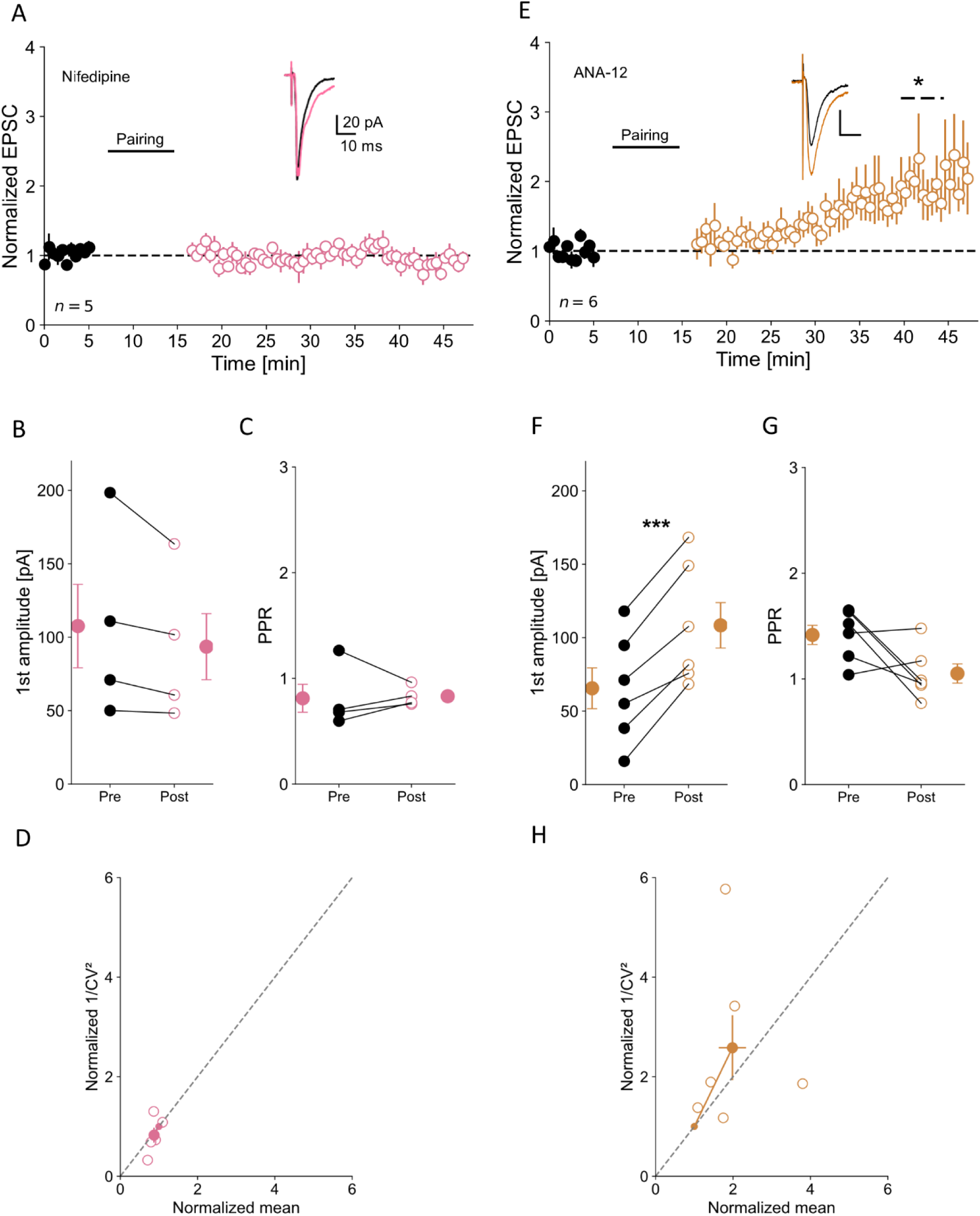
L-type VGCCs but not BDNF mediate synaptic plasticity at L2/3 VIP-IN excitatory inputs. **A)** Normalized EPSC after L-type VGCC blocker (nifedipine) application (pre pairing – black circles, post pairing – pink circles). Inset: representative EPSCs for pre and post LTP induction. **B)** Averaged 1^st^ amplitude for pre and post pairing protocol with blocked L-type VGCC activity. **C)** Averaged paired pulse ratio for pre and post pairing in the presence of nifedipine. **D)** 1/CV^2^ analysis in nifedipine. **E)** Same as A) but in the presence of the TrkB antagonist (ANA-12, pre pairing – black circles, post pairing – brown circles). **F-H)** Same as B-D) but in presence of ANA-12. Mean ± SEM, * p < 0.05.

Together, these findings suggest that postsynaptic mGluR-1 but not mGluR-5 is necessary for LTP of excitatory transmission to L2/3 VIP-INs. LTP induction depends on calcium influx through L-type VGCC but not TRPC.

### 3.4. Presynaptic expression of LTP in VIP-INs

Since PPR and 1/CV^2^ analyses (Fig. 1E, 2D,H, 4H) showed putative presynaptic locus of LTP expression, we next aimed to find a signaling factor acting retrogradely (Nicholson & Kullmann, 2014).

First, we checked whether brain-derived neurotrophic factor (BDNF) could be a retrograde messenger to trigger presynaptic changes in VIP-IN excitatory inputs, as it has been shown in CA1 neurons (Thapliyal *et al*., 2022) and cortical pyramidal neurons (Maglio *et al*., 2018). To do this, we blocked tropomyosin receptor kinase B (TrkB), a high-affinity receptor for BDNF, before the LTP pairing protocol. The TrkB receptor antagonist (ANA-12) did not prevent LTP induction (LTP was induced in 5 out of 6 neurons, 83 % LTP success rate, Fig. 5E, normalized amplitude 1.97 ± 0.34, p = 0.047, n = 6, one sample t-test; Fig. 5F, amplitude pre 65.57 ± 13.89 pA vs. post 108.42 ± 15.47 pA, p < 0.001, n = 6, paired t-test), indicating that BDNF is not necessary for LTP at excitatory inputs to L2/3 VIP-INs. Just like in our previous experiments, the PPR (Fig. 5G, pre 1.42 ± 0.09 vs. post 1.05 ± 0.09, p = 0.072, n = 6, paired t-test) and 1/CV^2^ analyses (Fig. 5H) revealed the presynaptic locus for LTP expression.

Next, we tested the involvement of the eCB retrograde signaling pathway in LTP. The release of eCBs is sensitive to extensive membrane depolarization and requires calcium influx that can be triggered by mGluR class I activation in this case (Heifets & Castillo, 2009). The eCB retrograde signaling pathway has been established to be an important factor in synaptic plasticity (Wang *et al*., 2016; Silva-Cruz *et al*., 2017; Maglio *et al*., 2018; Winters *et al*., 2023), however mainly implicated in LTD induction (Sjöström *et al*., 2003; Heifets & Castillo, 2009; Min & Nevian, 2012; Péterfi *et al*., 2012). Indeed, administration of the CBR-1 antagonist (AM-251) surprisingly blocked LTP in cortical VIP-INs. In this case, LTP was induced in 3 out of 13 cells (23 % LTP success rate, Fig. 6A, normalized amplitude 1.03 ± 0.11, p = 0.795, n = 13, one sample t-test; Fig. 6B, amplitude pre 64.71 ± 8.64 pA vs. post 69.49 ± 9.80 pA, p= 0.325, n = 13, paired t-test; Fig. 6C PPR pre 1.08 ± 0.11 vs. post 0.98 ± 0.07, p = 0.549, n = 12, paired t-test; Fig. 6D), suggesting that CBR-1 is responsible for presynaptic LTP expression.

**Fig. 6.**
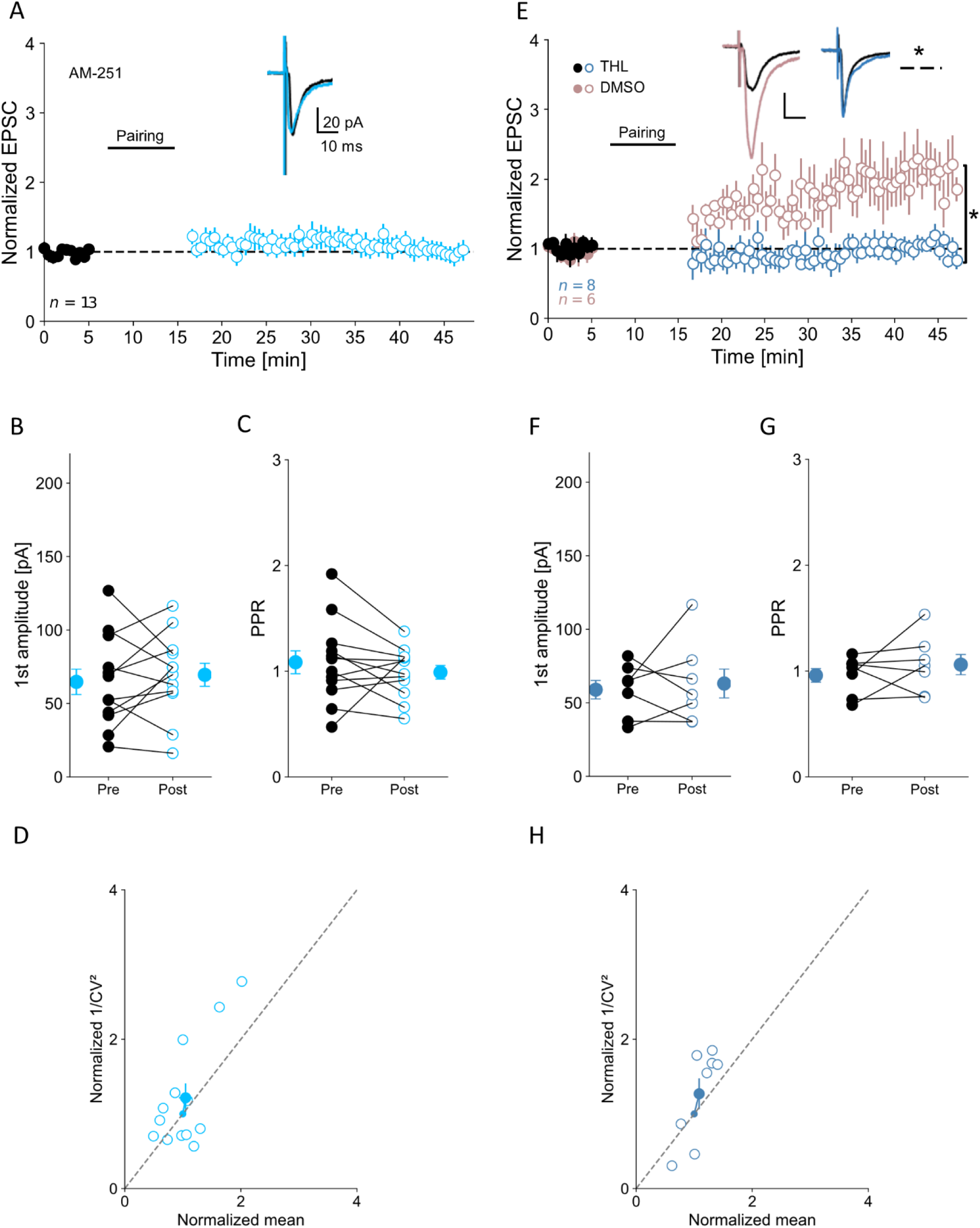
Postsynaptically synthesized eCBs putatively act retrogradely on glutamatergic inputs to VIP-INs promoting presynaptic expression of LTP. A) Time course plot of normalized. EPSC before and after the pairing protocol in the condition of eCBs signaling blockage with the CBR-1 antagonist (AM-251, pre pairing – black circles, post pairing – blue circles). Inset: EPSC traces for pre and post LTP induction. **B)** Averaged 1^st^ amplitude for pre and post pairing protocol in AM-251. **C)** Averaged paired pulse ratio for pre and post pairing in AM-251. **D)** 1/CV^2^ analysis in AM-251 conditions. **E)** Same as A) but in the presence of the lipase inhibitor (THL) inhibiting eCB synthesis (pre pairing – black circles, post pairing – navy blue circles) and with vehicle (DMSO, pre pairing – solid beige circles, post pairing – open beige circles). Horizontal dashed line with an asterisk indicates statistically significant differences for LTP induction in the vehicle control, vertical line with an asterisk indicates a significant difference between two recording conditions. Inset: examples of EPSC traces for pre and post LTP induction in both conditions. **F-H)** Same as B-D) but with THL. Mean ± SEM, * p < 0.05.

To verify whether a source of eCBs was postsynaptic, we filled VIP-INs with internal solution containing a lipase inhibitor (THL), to prevent eCB synthesis in the postsynaptic cell. In this condition, LTP was abolished in 5 out 8 cells (38 % LTP success rate, Fig. 6E, normalized amplitude 1.05 ± 0.09, p = 0.635, n = 8, one sample t-test; Fig. 6F, amplitude pre 58.92 ± 6.29 pA vs. post 63.13 ± 9.80 pA, p = 0.689, n = 8, paired t-test; Fig. 6G, PPR pre 0.96 ± 0.06 vs. post 1.06 ± 0.09, p = 0.376, n = 8, paired t-test). Additionally, we performed control experiments with a vehicle (DMSO in internal solution) showing that DMSO has no effect on LTP induction in 5 out of 6 cells (83 % LTP success rate, Fig. 6E, normalized amplitude 2.08 ± 0.33, p = 0.021, n = 6, one sample t-test; THL vs. DMSO p = 0.005, independent t-test).

Finally, we tested a possibility of adenosine-mediated signaling in LTP at VIP-INs (Falcón-Moya *et al*., 2020; Martínez-Gallego *et al*., 2022). For this purpose, the adenosine type 2 receptor (A2R) antagonist (SCH58261) was bath applied. A2R blockage failed to prevent LTP induction at excitatory inputs to L2/3 VIP-INs. In this condition, LTP was induced in 6 out of 7 neurons (86 % success rate, Fig. 7A, normalized amplitude 2.49 ± 0.48, p = 0.03, n = 7, one sample t-test; Fig. 7B, amplitude pre 47.29 ± 3.78 pA vs. post 96.11 ± 15.14 pA, p = 0.034, n = 7, paired t-test; Fig. 7C, PPR pre 1.31 ± 0.09 vs. post 1.02 ± 0.08, p = 0.016, n = 7, Wilcoxon test). However, when adenosine type 1 receptor (A1R) antagonist (DPCPX) was administered, LTP was diminished and LTP success rate dropped to 40 % (LTP was induced successfully in 2 out 5 cells, Fig. 7E, normalized amplitude 0.97 ± 0.21, p = 0.790, n = 5, one sample t-test; Fig. 7F, amplitude pre 58.32 ± 7.95 pA vs. post 54.06 ± 12.16 pA, p = 0.756, n = 5, paired t-test; Fig. 7G, PPR pre 0.91 ± 0.08 vs. post 0.96 ± 0.11, p = 0.721, n = 5, paired t-test).

**Fig. 7.**
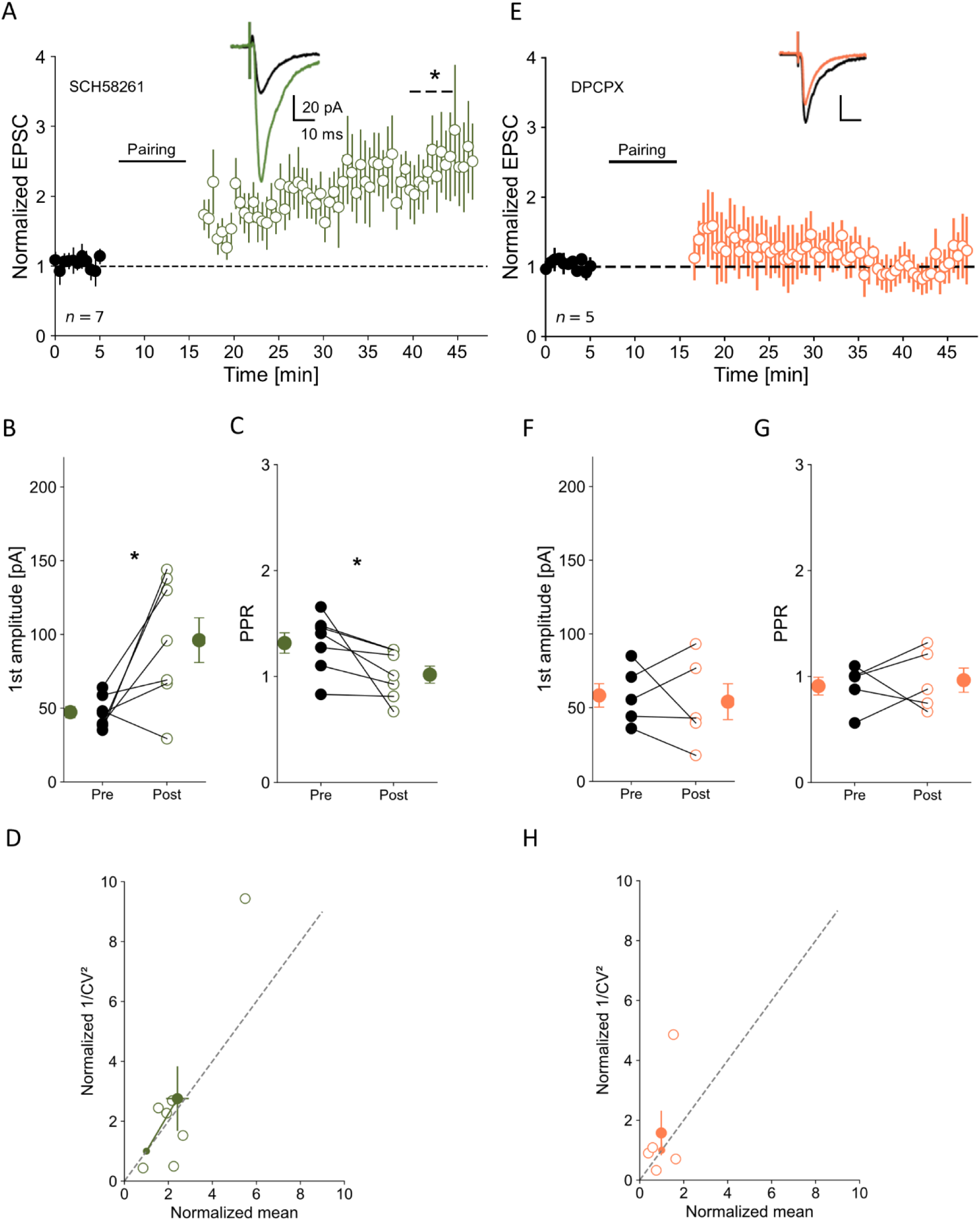
A1R but not A2R mediates LTP expression. **A)** Normalized EPSC before and after the pairing protocol in the presence of A2R antagonist (SCH58261: pre pairing – black circles, post pairing – green circles) Inset: examples of EPSC traces for pre and post LTP induction. **B)** Averaged 1^st^ amplitude for pre and post pairing protocol in SCH58261. **C)** Averaged paired pulse ratio for pre and post pairing in SCH58261. **D)** 1/CV^2^ analysis in A2R blockage conditions. **E)** Same as A) but in the presence of A1Rs antagonist (DPCPX) (pre pairing – black circles, post pairing – orange circles). **F-H)** Same as B-D) but in DPCPX bath-applied. Mean ± SEM, * p < 0.05.

In conclusion, our results indicate that expression of LTP at excitatory connections to L2/3 VIP-INs depends on retrograde signaling mediated by the postsynaptic release of eCBs, but not BDNF. Additionally, A1Rs are involved in LTP, suggesting astrocyte-dependent LTP at VIP-INs.

## 4. Discussion

Our findings provide insight into the mechanism of the Hebbian long-term plasticity of excitatory synapses on VIP interneurons in L2/3 of the mouse somatosensory cortex. Here, using electrophysiological approach, we described molecular factors essential for LTP induction and expression in these interneurons. We established that LTP at excitatory inputs to VIP-INs is mediated by postsynaptic activation of mGluR-1 and L-type VGCC. Moreover, our findings uncovered CBR-1 as the presynaptic target of LTP expression at excitatory synapses on VIP-INs, revealing a non-canonical function of eCBs in synaptic plasticity. Additionally, our data suggest the involvement of astrocytes in LTP at these connections.

It has been hypothesized that LTP cannot occur at excitatory inputs to GABAergic neurons (McBain *et al*., 1999; Ross & Soltesz, 2001), nevertheless later studies have shown that plasticity direction is brain region- and synapse type-specific (Perez *et al*., 2001; Topolnik *et al*., 2006; Sarihi *et al*., 2008; Oren *et al*., 2009; Banerjee *et al*., 2009; Chen *et al*., 2009; Szegedi *et al*., 2016; Chistiakova *et al*., 2019; McFarlan *et al*., 2024b). Similarly to research conducted on excitatory inputs to L2/3 VIP-INs in the motor cortex (McFarlan *et al*., 2024b) and on hippocampal CA1 inputs to SST-INs (Vasuta *et al*., 2015), we were able to induce LTP at excitatory synapses on L2/3 VIP-INs in the barrel cortex, even with no pharmacological blockade of GABAaRs. It is worthwhile to mention that synaptic plasticity is an interplay of glutamatergic and GABAergic net force (Wang & Maffei, 2014; Field *et al*., 2020; Agnes & Vogels, 2024). Many studies have shown that across different brain regions, GABAaRs can either block or conversely, facilitate LTP induction (Ruiz *et al*., 2010; Paille *et al*., 2013; Davenport *et al*., 2021). Indeed, pharmacological blockade of GABAaRs enhanced LTP at excitatory synapses on L2/3 VIP-INs in the barrel cortex, inferring that in physiological conditions those receptors diminish LTP at these connections. It has been suggested that blocking GABAaR-mediated inhibition facilitates postsynaptic membrane depolarization and thus enhances activation of NMDARs or voltage-dependent channels during LTP induction (Grover & Yan, 1999). Since in our study, the postsynaptic neurons were artificially depolarized during the LTP protocol, we assume that release of presynaptic neurons from GABAaR inhibition might have essential impact in facilitation of LTP in this case. Alternatively, there might be a more technical explanation of how blocking GABAaRs facilitates LTP. Namely, pharmacological blockade of GABAaRs reduces membrane conductance leading to more effective whole-cell clamping thus allowing detection of larger EPSCs.

Numerous studies have recognized NMDAR as the primary glutamatergic signaling factor of synaptic plasticity in the central nervous system, especially in the hippocampus (Citri & Malenka, 2008; Hirai *et al*., 2022), but also in thalamocortical inputs to L4 (Lu *et al*., 2001), in excitatory connections to pyramidal neurons located in L2/3 (Banerjee *et al*., 2009, 2014; Williams & Holtmaat, 2019) and L5 of the somatosensory cortex (Maglio *et al*., 2018; Gómez *et al*., 2021) or in motor cortex (Hasan *et al*., 2013). NMDARs are vastly expressed in VIP-INs (Porter *et al*., 1998).

The pairing protocol applied to induce plasticity in VIP-INs is highly dependent on postsynaptic calcium signaling triggered by membrane depolarization, favouring the NMDAR activation (Feldman, 2000). Therefore, we expected NMDAR to be a key player necessary for LTP induction in L2/3 VIP-INs of the mouse somatosensory cortex. To our surprise, induction of LTP at glutamatergic synapses on VIP-INs was NMDAR-independent similarly to what has been observed in another class of inhibitory interneurons – SST-INs in the hippocampus and neocortex (Perez *et al*., 2001; Oren *et al*., 2009; Chen *et al*., 2009; Nicholson & Kullmann, 2014). In contrast, LTP is NMDAR-dependent at excitatory inputs to hippocampal parvalbumin interneurons (PV-INs) (Cornford *et al*., 2019) or between cortical pyramidal cells (Banerjee *et al*., 2009).

Other receptors underlying plasticity formation are mGluRs or ionotropic receptors such as calcium-permeable AMPARs and nicotinic receptors (Alkadhi, 2021; McFarlan *et al*., 2023). In the present study, we assessed that mGluR-5 does not contribute to LTP at excitatory inputs to L2/3 VIP-INs, unlike at glutamatergic inputs to L2/3 pyramidal neurons (Cui *et al*., 2018). However, mGluR-1 is necessary for LTP induction in L2/3 VIP-INs, as it has been observed for excitatory synapses on SST-INs in hippocampal CA1 oriens/alveus (Vasuta *et al*., 2015). PV-INs, in turns, show mostly mGluR-5-dependent plasticity, possibly since there are hardly any mGluR-1 expressed in those neurons (Sarihi *et al*., 2008; Vasuta *et al*., 2015). Interestingly, in the human cortex, mGluR class I has been implicated in LTD induction in pyramidal to pyramidal connections (Kroon *et al*., 2019), as well as in excitatory inputs to fast-spiking interneurons (putative PV-INs) (Szegedi *et al*., 2016). Moreover, there are neurons that require activity of both mGluR-1 and −5 for LTP induction, as it has been observed in corticostriatal synaptic connections (Gubellini *et al*., 2003).

Metabotropic receptors regulate an increase of the calcium ion concentration in the postsynaptic cell via different signaling pathways (Mannaioni *et al*., 2001; Topolnik *et al*., 2006; Eng *et al*., 2016). It has been previously discovered that mGluR-1 can mediate synaptic plasticity in CA1 via the MAPK cascade (Topolnik *et al*., 2006; Eng *et al*., 2016). Topolnik et al. (Topolnik *et al*., 2006) have shown that hippocampal O/A interneurons require the Src/ERK downstream pathway in order to activate MAPK and thus induce LTP. On the contrary, we uncovered that LTP at glutamatergic inputs to cortical VIP-INs are dependent solely on Src tyrosine kinase activity, since blockage of this enzyme abolishes LTP completely, whereas in the hippocampus, LTP required complementary activity of ERK (Topolnik *et al*., 2006). Additionally, Src activation has been established to lead to calcium ion influx through TRPC (Topolnik *et al*., 2006; Kubota *et al*., 2014) but our data excluded TRPC involvement in LTP induction in VIP-INs. However, we found that LTP in these interneurons is depended on L-type VGCCs, suggesting that depolarization during the pairing protocol profoundly activates these channels leading to calcium influx necessary for LTP induction. L-type VGCC- but not NMDA-dependent LTP has previously been observed in many brain areas including the somatosensory cortex, amygdala and CA1 (Grover & Teyler, 1990; Weisskopf *et al*., 1999; Cui *et al*., 2018). Nonetheless, it remains obscure whether Src kinase directly interacts with L-type VGCC in VIP-INs. Previous single-channel recordings of hippocampal neurons have revealed that Src can directly boost L-type VGCC activity (Man *et al*., 2023). Also, studies on neuronal L-type VGCC expressed in HEK293-T showed physical binding of Src to these channels (Chao *et al*., 2011). In light of these two lines of evidence, we may hypothesize that Src kinase binds to and activates L-type VGCCs, or more likely the enzyme enhances the activity of these channels which are already activated through depolarization of the postsynaptic membrane during the paring protocol. Also, we cannot rule out involvements of presynaptic L-type VGCCs as well as astrocytic channels in the process of LTP induction (Falcón-Moya *et al*., 2020). However, we established that LTP at excitatory inputs to L2/3 VIP-INs requires postsynaptic G protein signaling, presumably triggered by mGluR-1 activation.

Synaptic plasticity may arise from changes in the postsynaptic or presynaptic membrane or be a result of mixed effects – pre- and postsynaptic mechanisms. It has been found that a direction of plasticity is dependent on the age and stimulation location (Reyes & Sakmann, 1999; Banerjee *et al*., 2009, 2014; St. Laurent & Kauer, 2019). Here, even though induction of LTP in VIP-INs is postsynaptic, analyses of PPRs and 1/CV^2^ suggest that LTP expression locus lays in the presynaptic site. Corresponding results based on these analyses have been observed in cortical excitatory inputs onto SST-INs (Chen *et al*., 2009; Nicholson & Kullmann, 2014; Chistiakova *et al*., 2019) and in excitatory synapses on hippocampal oriens-lacunosum moleculare (O-LM) interneurons (Oren *et al*., 2009). On the contrary, postsynaptic LTP expression has been found at glutamatergic inputs to L2/3 VIP-INs in the motor cortex (McFarlan *et al*., 2024b). The discrepancies between our results and McFarlan’s observation may result from the following factors: diverse brain regions (somatosensory cortex vs. motor cortex), age (P21-28 vs. P21-40), stimulation protocols (pairing postsynaptic depolarization with external stimulation vs. spike timing-dependent plasticity) and the location of stimulation electrodes (below patched cells vs. above), respectively.

Surprisingly, we found that LTP at excitatory inputs onto L2/3 VIP-INs is depended on CBR-1 activity, suggesting a role of eCBs as retrograde messengers in this process. Canonically, activation of CBR-1s reduces neurotransmitter release, and thus contributes to a decrease in synaptic transmission (Sjöström *et al*., 2003; Chevaleyre & Castillo, 2004; Chevaleyre *et al*., 2006; Heifets & Castillo, 2009; Min & Nevian, 2012; Péterfi *et al*., 2012). Nonetheless, growing body of evidence also indicates contribution of eCBs to LTP (Wang *et al*., 2016; Silva-Cruz *et al*., 2017; Maglio *et al*., 2018; Cui *et al*., 2018; Winters *et al*., 2023). Similarly to our findings, eCBs have been reported to mediate LTP in L5 and 2/3 pyramidal neurons in the rat barrel cortex (Maglio *et al*., 2018; Cui *et al*., 2018). Furthermore, Maglio et al. study showed that eCB-dependent LTP in pyramidal neurons also relays on BNDF, in contrast to our findings, in which TrkB activation was not crucial for LTP in VIP-INs. The location of CBR-1s contributing to LTP at excitatory connections to VIP-INs, remains undiscovered. These receptors could be located on both the presynaptic site and astrocytes (Sjöström *et al*., 2003; Navarrete & Araque, 2010; Min & Nevian, 2012; Wang *et al*., 2016; Martin-Fernandez *et al*., 2017; Falcón-Moya *et al*., 2020; Martínez-Gallego *et al*., 2022; Winters *et al*., 2023). Since blocking A1Rs prevents LTP induction, astrocytes might be involved in this process. As depicted in Fig. 8, eCBs released postsynaptically can activate CB1R on astrocyte, leading to a signaling pathway that triggers the release of astrocytic adenosine that in turn binds to A1R on the presynaptic site (Falcón-Moya *et al*., 2020; Martínez-Gallego *et al*., 2022). In another scenario, both CB1R and A1R could be located presynaptically, and LTP is induced with or without astrocyte involvement (Ohno-Shosaku *et al*., 2012; Sitzia *et al*., 2023; Nasrallah *et al*., 2024). We hypothesize that VIP-IN depolarization maintained during the LTP protocol leads to the eCB release from the postsynaptic site, because it has been established that eCBs are synthesized in an activity-dependent manner during an intense membrane depolarization (Di Marzo *et al*., 1998; Heifets & Castillo, 2009). Our experiments showed that the lipase inhibitor loaded to the postsynaptic cell indeed blocks LTP at excitatory synapses on VIP-INs, indicating the postsynaptic source of eCBs for LTP induction. Interestingly, the age of an animal might determine the contribution of eCBs to either LTD or LTP, as demonstrated for a developmental profile of NMDAR-dependent spike-timing plasticity (Banerjee *et al*., 2009). Namely, eCBs have been implicated in LTD at excitatory inputs from L4 to L2/3 pyramidal cells of the somatosensory cortex of young rats (16-21 day old) (Min & Nevian, 2012). However, in older rats (25-35 day old), eCBs elicited LTP at these connections (Cui *et al*., 2018). It would be worthwhile to test this hypothesis in younger mice as well (e.g. 14-18 day old). Interestingly, eCB-LTP or eCB-LTD can be evoked in the same neuron sequentially by using appropriate spike timing-dependent protocols (Cui *et al*., 2016). What’s more, in Min and Nevian study, eCB-LTD also was mediated by NMDARs (Min & Nevian, 2012), however our findings support Cui et al. results, in which eCB-LTP was NMDAR-independent (Cui *et al*., 2018). Nonetheless, eCB involvement in the presynaptic LTP remains obscure. One explanation could be that CBR-1 is implicated in LTP, as in the striatum and neocortex, due to an interplay with the transient receptor potential vanilloid-1 (TRPV1) (Cui *et al*., 2015, 2018). As TRPV1 channels are highly permeable to calcium (Ross, 2003), they may contribute to LTP by boosting calcium transient in the presynaptic site which in turn, triggers an increase of neurotransmitter release from excitatory terminals on L2/3 VIP-INs. Another possibility could be that G proteins coupled to CB1Rs trigger a signaling pathway acting on internal calcium stores to elevate calcium levels in the cytoplasm. Exact mechanisms of CB1R action in LTP remain yet to be experimentally dissected.

**Fig. 8.**
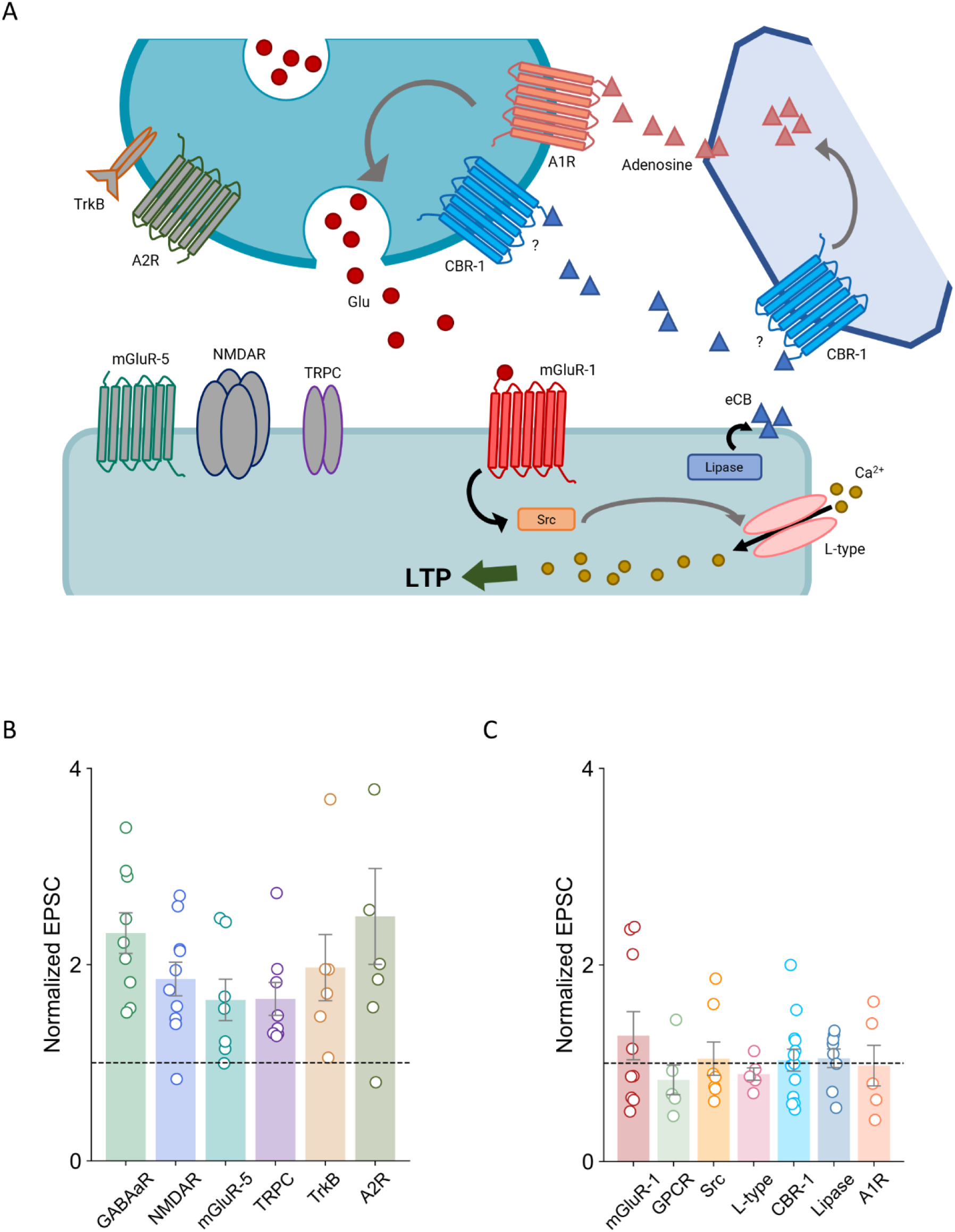
Summary of factors tested for the contribution in LTP at excitatory inputs to L2/3 VIP-INs in the mouse somatosensory cortex. **A)** Schematic representation of a glutamatergic synapse on a VIP-IN. During LTP protocol, glutamate binding to the postsynaptic mGluR-1 presumably activates Src-kinase signaling cascade necessary for LTP induction. Additionally, membrane depolarization during LTP protocol, activates L-type VGCC leading to the postsynaptic calcium influx needed for LTP induction. Postsynaptically released eCBs act as retrograde messengers, activating CBR-1s on either the presynaptic site or an astrocyte or both, leading to the presynaptic LTP expression. Also, A1R activation is involved in LTP. NMDAR, mGluR-5, TRPC, BDNF (TrkB) and A2R do not mediate LTP in VIP-INs (receptors to the left, showed in gray). **B)** Summarized graph showing receptors and channels that are not necessary for LTP. Pharmacological interventions did not abolish LTP. **C)** Summarized plot depicting crucial factors for synaptic plasticity at excitatory synapses on VIP-INs. Here, pharmacological agents blocked LTP.

Finally, nitric oxide (NO) has been shown to play a crucial role as a retrograde messenger in hippocampal and neocortical LTP (Hardingham & Fox, 2006; Padamsey *et al*., 2017; Falcón-Moya *et al*., 2020). However, contrary to hippocampal VIP-INs (Tricoire *et al*., 2010), neocortical VIP-INs rather scarcely (only up to 10 % of the population) express NO synthetase (Karagiannis *et al*., 2009; Perrenoud *et al*., 2012). Hence, NO-mediated signaling in LTP at excitatory synapses onto L2/3 VIP-INs is unlikely involved.

What is a consequence of LTP at excitatory connections to L2/3 VIP-INs in the barrel cortex? Enhanced excitatory transmission to VIP-INs might serve as a positive feedback for stronger VIP-IN-mediated inhibition of SST-INs, leading to disinhibition of excitatory neurons from SST-IN-mediated inhibition during learning process. On the other hand, greater excitation of VIP-INs might be a part of a pathological mechanism leading to network hyperexcitation, because it has been found that optogenetic and genetic silencing of VIP-INs reduces susceptibility to seizures (Khoshkhoo *et al*., 2017; Rahimi *et al*., 2023). In this case, the LTP pairing protocol with prolonged VIP-IN depolarization might serve as a model to study mechanisms of pathological conditions.

## 5. Acknowledgments

This work was supported by the National Science Centre, Poland (2020/39/B/NZ4/01462) to JUC.

## 8. Data Availability Statement

Data is available on OSF (https://osf.io/user/gw85z).

## 9. Author Contributions

Karolina Bogaj: conceptualization, methodology, investigation, data curation, formal analysis, visualization, writing – original draft, writing – review and editing. Joanna Urban-Ciecko: conceptualization, methodology, data curation, supervision, funding acquisition, writing – review and editing.

## 10. Abbreviations

ACSF: artificial cerebrospinal fluid
AM-251: N-(Piperidin-1-yl)-5-(4-iodophenyl)-1-(2,4-dichlorophenyl)-4-methyl-1H-pyrazole-3-carboxamide
ANA-12: N-[2-[[(Hexahydro-2-oxo-1H-azepin-3-yl)amino]carbonyl]phenyl]benzo[b]thiophene-2-carboxamide
APV: D-(-)-2-Amino-5-phosphonopentanoic acid
A1R: adenosine type 1 receptor
A2R: adenosine type 2 receptor
BDNF: brain-derived neurotrophic factor
CBR-1: cannabinoid receptor 1
CP-AMPAR: calcium permeable-α-amino-3-hydroxy-5-methyl-4-isoxazolepropionic acid receptor
CV: coefficient of variation
DMSO: dimethyl sulfoxide
DPCPX: 8-Cyclopentyl-1,3-dipropylxanthine
eCBs: endocannabinoids
EPSCs: excitatory postsynaptic currents
GABAaR: γ-aminobutyric acid type A receptor
GDPβS: Guanosine-5’-(β-thio)-diphosphate
GPCR: G protein-coupled receptor
L-type VGCC: long-lasting voltage-gated calcium channel
LTD: long-term depression
LTP: long-term potentiation
LY367385: (S)-(+)-α-Amino-4-carboxy-2-methylbenzeneacetic acid
mGluR: metabotropic glutamate receptor
MTEP: 3-((2-Methyl-1,3-thiazol-4-yl)ethynyl)pyridine hydrochloride
NMDAR: N-methyl-D-aspartate receptor
PKC: protein kinase C
PPR: paired pulse ratio
PP2: 3-(4-chlorophenyl) 1-(1,1-dimethylethyl)-1H-pyrazolo[3,4-d]pyrimidin-4-amine
SCH58261: 2-(2-Furanyl)-7-(2-phenylethyl)-7H-pyrazolo[4,3-e][1,2,4]triazolo[1,5-c]pyrimidin-5-amine
SKF96365: 1-[2-(4-Methoxyphenyl)-2-[3-(4-methoxyphenyl)propoxy]ethyl-1H-imidazole hydrochloride
THL: tetrahydrolipstatin
TrkB: tropomyosin receptor kinase B
TRPC: transient receptor potential channel
VIP-INs: vasoactive intestinal polypeptide-expressing interneurons

## References

Agnes EJ & Vogels TP (2024). Co-dependent excitatory and inhibitory plasticity accounts for quick, stable and long-lasting memories in biological networks. Nat Neurosci 27, 964–974.

Alkadhi KA (2021). NMDA receptor-independent LTP in mammalian nervous system. Progress in Neurobiology 200, 101986.

Audette NJ, Urban-Ciecko J, Matsushita M & Barth AL (2018). POm Thalamocortical Input Drives Layer-Specific Microcircuits in Somatosensory Cortex. Cerebral Cortex 28, 1312–1328.

Banerjee A, González-Rueda A, Sampaio-Baptista C, Paulsen O & Rodríguez-Moreno A (2014). Distinct mechanisms of spike timing-dependent LTD at vertical and horizontal inputs onto L2/3 pyramidal neurons in mouse barrel cortex. Physiol Rep 2, e00271.

Banerjee A, Meredith RM, Rodríguez-Moreno A, Mierau SB, Auberson YP & Paulsen O (2009). Double Dissociation of Spike Timing–Dependent Potentiation and Depression by Subunit-Preferring NMDA Receptor Antagonists in Mouse Barrel Cortex. Cerebral Cortex 19, 2959–2969.

Bender VA, Pugh JR & Jahr CE (2009). Presynaptically Expressed Long-Term Potentiation Increases Multivesicular Release at Parallel Fiber Synapses. J Neurosci 29, 10974–10978.

Benquet P, Gee CE & Gerber U (2002). Two Distinct Signaling Pathways Upregulate NMDA Receptor Responses via Two Distinct Metabotropic Glutamate Receptor Subtypes. J Neurosci 22, 9679–9686.

Bogaj K, Kaplon R & Urban-Ciecko J (2023). GABAAR-mediated tonic inhibition differentially modulates intrinsic excitability of VIP- and SST- expressing interneurons in layers 2/3 of the somatosensory cortex. Front Cell Neurosci 17, 1270219.

Bogaj K & Urban-Ciecko J (2025). Inhibition of BK channels by GABAb receptors enhances intrinsic excitability of layer 2/3 vasoactive intestinal polypeptide-expressing interneurons in mouse neocortex. The Journal of Physiology 603, 1171–1196.

Booker SA, Loreth D, Gee AL, Watanabe M, Kind PC, Wyllie DJA, Kulik Á & Vida I (2018). Postsynaptic GABABRs Inhibit L-Type Calcium Channels and Abolish Long-Term Potentiation in Hippocampal Somatostatin Interneurons. Cell Reports 22, 36–43.

Bowie D & Mayer ML (1995). Inward rectification of both AMPA and kainate subtype glutamate receptors generated by polyamine-mediated ion channel block. Neuron 15, 453–462.

Brock JA, Thomazeau A, Watanabe A, Li SSY & Sjöström PJ (2020). A Practical Guide to Using CV Analysis for Determining the Locus of Synaptic Plasticity. Front Synaptic Neurosci 12, 11.

Canto-Bustos M, Friason FK, Bassi C & Oswald A-MM (2022). Disinhibitory Circuitry Gates Associative Synaptic Plasticity in Olfactory Cortex. J Neurosci 42, 2942–2950.

Chao J-T, Gui P, Zamponi GW, Davis GE & Davis MJ (2011). Spatial association of the Cav1.2 calcium channel with α_5_β_1_-integrin. American Journal of Physiology-Cell Physiology 300, C477–C489.

Chavis P, Ango F, Michel J, Bockaert J & Fagni L (1998). Modulation of big K^+^ channel activity by ryanodine receptors and L-type Ca^2+^ channels in neurons. Eur J of Neuroscience 10, 2322–2327.

Chavis P, Fagni L, Lansman JB & Bockaert J (1996). Functional coupling between ryanodine receptors and L-type calcium channels in neurons. Nature 382, 719–722.

Chen H-X, Jiang M, Akakin D & Roper SN (2009). Long-term potentiation of excitatory synapses on neocortical somatostatin-expressing interneurons. J Neurophysiol 102, 3251–3259.

Chéreau R, Williams LE, Bawa T & Holtmaat A (2022). Circuit mechanisms for cortical plasticity and learning. Seminars in Cell & Developmental Biology 125, 68–75.

Chevaleyre V & Castillo PE (2004). Endocannabinoid-Mediated Metaplasticity in the Hippocampus. Neuron 43, 871–881.

Chevaleyre V, Takahashi KA & Castillo PE (2006). ENDOCANNABINOID-MEDIATED SYNAPTIC PLASTICITY IN THE CNS. Annu Rev Neurosci 29, 37–76.

Chistiakova M, Ilin V, Roshchin M, Bannon N, Malyshev A, Kisvárday Z & Volgushev M (2019). Distinct Heterosynaptic Plasticity in Fast Spiking and Non-Fast-Spiking Inhibitory Neurons in Rat Visual Cortex. J Neurosci 39, 6865–6878.

Citri A & Malenka RC (2008). Synaptic Plasticity: Multiple Forms, Functions, and Mechanisms. Neuropsychopharmacol 33, 18–41.

Cornford JH, Mercier MS, Leite M, Magloire V, Häusser M & Kullmann DM (2019). Dendritic NMDA receptors in parvalbumin neurons enable strong and stable neuronal assemblies. eLife 8, e49872.

Cui Y, Paillé V, Xu H, Genet S, Delord B, Fino E, Berry H & Venance L (2015). Endocannabinoids mediate bidirectional striatal spike-timing-dependent plasticity. The Journal of Physiology 593, 2833–2849.

Cui Y, Perez S & Venance L (2018). Endocannabinoid-LTP Mediated by CB1 and TRPV1 Receptors Encodes for Limited Occurrences of Coincident Activity in Neocortex. Front Cell Neurosci; DOI: 10.3389/fncel.2018.00182.

Cui Y, Prokin I, Xu H, Delord B, Genet S, Venance L & Berry H (2016). Endocannabinoid dynamics gate spike-timing dependent depression and potentiation. eLife 5, e13185.

Davenport CM, Rajappa R, Katchan L, Taylor CR, Tsai M-C, Smith CM, De Jong JW, Arnold DB, Lammel S & Kramer RH (2021). Relocation of an Extrasynaptic GABAA Receptor to Inhibitory Synapses Freezes Excitatory Synaptic Strength and Preserves Memory. Neuron 109, 123–134.e4.

De León-López CAM, Carretero-Rey M & Khan ZU (2025). AMPA Receptors in Synaptic Plasticity, Memory Function, and Brain Diseases. Cell Mol Neurobiol 45, 14.

Di Marzo V, Melck D, Bisogno T & De Petrocellis L (1998). Endocannabinoids: endogenous cannabinoid receptor ligands with neuromodulatory action. Trends in Neurosciences 21, 521–528.

Eng AG, Kelver DA, Hedrick TP & Swanson GT (2016). Transduction of group I mGluR-mediated synaptic plasticity by β-arrestin2 signalling. Nat Commun 7, 13571.

Falcón-Moya R, Pérez-Rodríguez M, Prius-Mengual J, Andrade-Talavera Y, Arroyo-García LE, Pérez-Artés R, Mateos-Aparicio P, Guerra-Gomes S, Oliveira JF, Flores G & Rodríguez-Moreno A (2020). Astrocyte-mediated switch in spike timing-dependent plasticity during hippocampal development. Nat Commun 11, 4388.

Feldman DE (2000). Timing-Based LTP and LTD at Vertical Inputs to Layer II/III Pyramidal Cells in Rat Barrel Cortex. Neuron 27, 45–56.

Field RE, D’amour JA, Tremblay R, Miehl C, Rudy B, Gjorgjieva J & Froemke RC (2020). Heterosynaptic Plasticity Determines the Set Point for Cortical Excitatory-Inhibitory Balance. Neuron 106, 842–854.e4.

Fu Y, Kaneko M, Tang Y, Alvarez-Buylla A & Stryker MP (2015). A cortical disinhibitory circuit for enhancing adult plasticity. eLife; DOI: 10.7554/elife.05558.

Georgiou C, Kehayas V, Lee KS, Brandalise F, Sahlender DA, Blanc J, Knott G & Holtmaat A (2022). A subpopulation of cortical VIP-expressing interneurons with highly dynamic spines. Commun Biol 5, 352.

Gómez R, Maglio LE, Gonzalez-Hernandez AJ, Rivero-Pérez B, Bartolomé-Martín D & Giraldez T (2021). NMDA receptor–BK channel coupling regulates synaptic plasticity in the barrel cortex. Proc Natl Acad Sci USA 118, e2107026118.

Grover LM & Teyler TJ (1990). Two components of long-term potentiation induced by different patterns of afferent activation. Nature 347, 477–479.

Grover LM & Yan C (1999). Blockade of GABA_A_ Receptors Facilitates Induction of NMDA Receptor-Independent Long-Term Potentiation. Journal of Neurophysiology 81, 2814–2822.

Gubellini P, Saulle E, Centonze D, Costa C, Tropepi D, Bernardi G, Conquet F & Calabresi P (2003). Corticostriatal LTP requires combined mGluR1 and mGluR5 activation. Neuropharmacology 44, 8–16.

Hardingham N & Fox K (2006). The Role of Nitric Oxide and GluR1 in Presynaptic and Postsynaptic Components of Neocortical Potentiation. J Neurosci 26, 7395–7404.

Hasan MT, Hernández-González S, Dogbevia G, Treviño M, Bertocchi I, Gruart A & Delgado-García JM (2013). Role of motor cortex NMDA receptors in learning-dependent synaptic plasticity of behaving mice. Nat Commun 4, 2258.

Heidinger V, Manzerra P, Wang XQ, Strasser U, Yu S-P, Choi DW & Behrens MM (2002). Metabotropic Glutamate Receptor 1-Induced Upregulation of NMDA Receptor Current: Mediation through the Pyk2/Src-Family Kinase Pathway in Cortical Neurons. J Neurosci 22, 5452–5461.

Heifets BD & Castillo PE (2009). Endocannabinoid Signaling and Long-Term Synaptic Plasticity. Annu Rev Physiol 71, 283–306.

Hirai H, Sakaba T & Hashimotodani Y (2022). Subcortical glutamatergic inputs exhibit a Hebbian form of long-term potentiation in the dentate gyrus. Cell Reports 41, 111871.

Huemmeke M, Eysel UT & Mittmann T (2002). Metabotropic glutamate receptors mediate expression of LTP in slices of rat visual cortex. Eur J of Neuroscience 15, 1641–1645.

Jia Z, Lu Y, Henderson J, Taverna F, Romano C, Abramow-Newerly W, Wojtowicz JM & Roder J (1998). Selective abolition of the NMDA component of long-term potentiation in mice lacking mGluR5. Learn Mem 5, 331–343.

Jiang S-N, Cao J-W, Liu L-Y, Zhou Y, Shan G-Y, Fu Y-H, Shao Y-C & Yu Y-C (2023). *Sncg*, *Mybpc1*, and *Parm1* Classify subpopulations of VIP-expressing interneurons in layers 2/3 of the somatosensory cortex. Cerebral Cortex 33, 4293–4304.

Jiang X, Shen S, Cadwell CR, Berens P, Sinz F, Ecker AS, Patel S & Tolias AS (2015). Principles of connectivity among morphologically defined cell types in adult neocortex. Science 350, aac9462.

Kanigowski D, Bogaj K, Barth AL & Urban-Ciecko J (2023). Somatostatin-expressing interneurons modulate neocortical network through GABAb receptors in a synapse-specific manner. Sci Rep 13, 8780.

Kanigowski D & Urban-Ciecko J (2024). Conditioning and pseudoconditioning differently change intrinsic excitability of inhibitory interneurons in the neocortex. Cerebral Cortex; DOI: 10.1093/cercor/bhae109.

Kanigowski D & Urban-Ciecko J (2025). Conditional learning increases inhibition of layer 4 excitatory neurons by somatostatin- and parvalbumin-expressing interneurons in the barrel cortex. J Neural Transm; DOI: 10.1007/s00702-025-02949-5.

Karagiannis A, Gallopin T, Dávid C, Battaglia D, Geoffroy H, Rossier J, Hillman EMC, Staiger JF & Cauli B (2009). Classification of NPY-Expressing Neocortical Interneurons. J Neurosci 29, 3642–3659.

Kerner JA, Standaert DG, Penney JB, Young AB & Landwehrmeyer GB (1997). Expression of group one metabotropic glutamate receptor subunit mRNAs in neurochemically identified neurons in the rat neostriatum, neocortex, and hippocampus. Molecular Brain Research 48, 259–269.

Khoshkhoo S, Vogt D & Sohal VS (2017). Dynamic, Cell-Type-Specific Roles for GABAergic Interneurons in a Mouse Model of Optogenetically Inducible Seizures. Neuron 93, 291–298.

Kofuji P & Araque A (2021). G-Protein-Coupled Receptors in Astrocyte–Neuron Communication. Neuroscience 456, 71–84.

Kroon T, Dawitz J, Kramvis I, Anink J, Obermayer J, Verhoog MB, Wilbers R, Goriounova NA, Idema S, Baayen JC, Aronica E, Mansvelder HD & Meredith RM (2019). Group I mGluR-Mediated Activation of Martinotti Cells Inhibits Local Cortical Circuitry in Human Cortex. Front Cell Neurosci 13, 315.

Kubota H, Nagao S, Obata K & Hirono M (2014). mGluR1-Mediated Excitation of Cerebellar GABAergic Interneurons Requires Both G Protein-Dependent and Src–ERK1/2-Dependent Signaling Pathways ed. Obukhov AG. PLoS ONE 9, e106316.

Lee S, Kruglikov I, Huang ZJ, Fishell G & Rudy B (2013). A disinhibitory circuit mediates motor integration in the somatosensory cortex. Nat Neurosci 16, 1662–1670.

Lu H-C, Gonzalez E & Crair MC (2001). Barrel Cortex Critical Period Plasticity Is Independent of Changes in NMDA Receptor Subunit Composition. Neuron 32, 619–634.

Maffei A, Nelson SB & Turrigiano GG (2004). Selective reconfiguration of layer 4 visual cortical circuitry by visual deprivation. Nat Neurosci 7, 1353–1359.

Maglio LE, Noriega-Prieto JA, Maraver MJ & Fernández De Sevilla D (2018). Endocannabinoid-Dependent Long-Term Potentiation of Synaptic Transmission at Rat Barrel Cortex. Cerebral Cortex 28, 1568–1581.

Man KNM, Bartels P, Henderson PB, Kim K, Shi M, Zhang M, Ho S-Y, Nieves-Cintron M, Navedo MF, Horne MC & Hell JW (2023). α1-Adrenergic receptor-PKC-Pyk2-Src signaling boosts L-type Ca2+ channel CaV1.2 activity and long-term potentiation in rodents. Elife 12, e79648.

Mannaioni G, Marino MJ, Valenti O, Traynelis SF & Conn PJ (2001). Metabotropic Glutamate Receptors 1 and 5 Differentially Regulate CA1 Pyramidal Cell Function. J Neurosci 21, 5925–5934.

Martin SJ, Grimwood PD & Morris RGM (2000). Synaptic Plasticity and Memory: An Evaluation of the Hypothesis. Annu Rev Neurosci 23, 649–711.

Martínez-Gallego I, Pérez-Rodríguez M, Coatl-Cuaya H, Flores G & Rodríguez-Moreno A (2022). Adenosine and Astrocytes Determine the Developmental Dynamics of Spike Timing-Dependent Plasticity in the Somatosensory Cortex. J Neurosci 42, 6038–6052.

Martin-Fernandez M, Jamison S, Robin LM, Zhao Z, Martin ED, Aguilar J, Benneyworth MA, Marsicano G & Araque A (2017). Synapse-specific astrocyte gating of amygdala-related behavior. Nat Neurosci 20, 1540–1548.

Mayer ML, Westbrook GL & Guthrie PB (1984). Voltage-dependent block by Mg2+ of NMDA responses in spinal cord neurones. Nature 309, 261–263.

McBain CJ, Freund TF & Mody I (1999). Glutamatergic synapses onto hippocampal interneurons: precision timing without lasting plasticity. Trends in Neurosciences 22, 228–235.

McFarlan AR, Chou CYC, Watanabe A, Cherepacha N, Haddad M, Owens H & Sjöström PJ (2023). The plasticitome of cortical interneurons. Nat Rev Neurosci 24, 80–97.

McFarlan AR, Gomez I, Chou CYC, Alcolado A, Costa RP & Sjöström PJ (2024a). The short-term plasticity of VIP interneurons in motor cortex. Front Synaptic Neurosci 16, 1433977.

McFarlan AR, Guo C, Gomez I, Weinerman C, Liang TA & Sjöström PJ (2024b). The spike-timing-dependent plasticity of VIP interneurons in motor cortex. Front Cell Neurosci 18, 1389094.

Min R & Nevian T (2012). Astrocyte signaling controls spike timing–dependent depression at neocortical synapses. Nat Neurosci 15, 746–753.

Nahir B, Lindsly C & Frazier CJ (2010). mGluR-mediated and endocannabinoid-dependent long-term depression in the hilar region of the rat dentate gyrus. Neuropharmacology 58, 712–721.

Naskar S, Qi J, Pereira F, Gerfen CR & Lee S (2021). Cell-type-specific recruitment of GABAergic interneurons in the primary somatosensory cortex by long-range inputs. Cell Reports 34, 108774.

Nasrallah K, Berthoux C, Hashimotodani Y, Chávez AE, Gulfo MC, Luján R & Castillo PE (2024). Retrograde adenosine/A2A receptor signaling facilitates excitatory synaptic transmission and seizures. Cell Reports 43, 114382.

Navarrete M & Araque A (2010). Endocannabinoids Potentiate Synaptic Transmission through Stimulation of Astrocytes. Neuron 68, 113–126.

Nicholson E & Kullmann DM (2014). Long-term potentiation in hippocampal oriens interneurons: postsynaptic induction, presynaptic expression and evaluation of candidate retrograde factors. Phil Trans R Soc B 369, 20130133.

Nicoll RA (2003). Expression mechanisms underlying long-term potentiation: a postsynaptic view ed. Bliss TVP, Collingridge GL & Morris RGM. Phil Trans R Soc Lond B 358, 721–726.

Nicoll RA (2017). A Brief History of Long-Term Potentiation. Neuron 93, 281–290.

Nowak L, Bregestovski P, Ascher P, Herbet A & Prochiantz A (1984). Magnesium gates glutamate-activated channels in mouse central neurones. Nature 307, 462–465.

Ohno-Shosaku T, Tanimura A, Hashimotodani Y & Kano M (2012). Endocannabinoids and Retrograde Modulation of Synaptic Transmission. Neuroscientist 18, 119–132.

Oren I, Nissen W, Kullmann DM, Somogyi P & Lamsa KP (2009). Role of Ionotropic Glutamate Receptors in Long-Term Potentiation in Rat Hippocampal CA1 Oriens-Lacunosum Moleculare Interneurons. J Neurosci 29, 939–950.

Padamsey Z, Tong R & Emptage N (2017). Glutamate is required for depression but not potentiation of long-term presynaptic function. eLife 6, e29688.

Paille V, Fino E, Du K, Morera-Herreras T, Perez S, Kotaleski JH & Venance L (2013). GABAergic Circuits Control Spike-Timing-Dependent Plasticity. J Neurosci 33, 9353–9363.

Patton MH, Thomas KT, Bayazitov IT, Newman KD, Kurtz NB, Robinson CG, Ramirez CA, Trevisan AJ, Bikoff JB, Peters ST, Pruett-Miller SM, Jiang Y, Schild AB, Nityanandam A & Zakharenko SS (2024). Synaptic plasticity in human thalamocortical assembloids. Cell Reports 44, 114503.

Perez Y, Morin F & Lacaille J-C (2001). A hebbian form of long-term potentiation dependent on mGluR1a in hippocampal inhibitory interneurons. Proc Natl Acad Sci USA 98, 9401–9406.

Perrenoud Q, Geoffroy H, Gauthier B, Rancillac A, Alfonsi F, Kessaris N, Rossier J, Vitalis T & Gallopin T (2012). Characterization of Type I and Type II nNOS-Expressing Interneurons in the Barrel Cortex of Mouse. Front Neural Circuits; DOI: 10.3389/fncir.2012.00036.

Péterfi Z, Urbán GM, Papp OI, Németh B, Monyer H, Szabó G, Erdélyi F, Mackie K, Freund TF, Hájos N & Katona I (2012). Endocannabinoid-Mediated Long-Term Depression of Afferent Excitatory Synapses in Hippocampal Pyramidal Cells and GABAergic Interneurons. J Neurosci 32, 14448–14463.

Porter JT, Cauli B, Staiger JF, Lambolez B, Rossier J & Audinat E (1998). Properties of bipolar VIPergic interneurons and their excitation by pyramidal neurons in the rat neocortex. Eur J of Neuroscience 10, 3617–3628.

Prönneke A, Scheuer B, Wagener RJ, Möck M, Witte M & Staiger JF (2015). Characterizing VIP Neurons in the Barrel Cortex of VIPcre/tdTomato Mice Reveals Layer-Specific Differences. Cereb Cortex 25, 4854–4868.

Rahimi S, Salami P, Matulewicz P, Schmuck A, Bukovac A, Ramos-Prats A, Tasan RO & Drexel M (2023). The role of subicular VIP-expressing interneurons on seizure dynamics in the intrahippocampal kainic acid model of temporal lobe epilepsy. Experimental Neurology 370, 114580.

Reyes A & Sakmann B (1999). Developmental Switch in the Short-Term Modification of Unitary EPSPs Evoked in Layer 2/3 and Layer 5 Pyramidal Neurons of Rat Neocortex. J Neurosci 19, 3827–3835.

Ross RA (2003). Anandamide and vanilloid TRPV1 receptors. Br J Pharmacol 140, 790–801.

Ross ST & Soltesz I (2001). Long-term plasticity in interneurons of the dentate gyrus. Proc Natl Acad Sci USA 98, 8874–8879.

Ruiz A, Campanac E, Scott RS, Rusakov DA & Kullmann DM (2010). Presynaptic GABAA receptors enhance transmission and LTP induction at hippocampal mossy fiber synapses. Nat Neurosci 13, 431–438.

Sarihi A, Jiang B, Komaki A, Sohya K, Yanagawa Y & Tsumoto T (2008). Metabotropic Glutamate Receptor Type 5-Dependent Long-Term Potentiation of Excitatory Synapses on Fast-Spiking GABAergic Neurons in Mouse Visual Cortex. J Neurosci 28, 1224–1235.

Silva-Cruz A, Carlström M, Ribeiro JA & Sebastião AM (2017). Dual Influence of Endocannabinoids on Long-Term Potentiation of Synaptic Transmission. Front Pharmacol 8, 921.

Sitzia G, Abrahao KP, Liput D, Calandra GM & Lovinger DM (2023). Distinct mechanisms of CB1 and GABA_B_ receptor presynaptic modulation of striatal indirect pathway projections to mouse globus pallidus. The Journal of Physiology 601, 195–209.

Sjöström PJ, Turrigiano GG & Nelson SB (2003). Neocortical LTD via Coincident Activation of Presynaptic NMDA and Cannabinoid Receptors. Neuron 39, 641–654.

St. Laurent R & Kauer J (2019). Synaptic Plasticity at Inhibitory Synapses in the Ventral Tegmental Area Depends upon Stimulation Site. eNeuro 6, ENEURO.0137-19.2019.

Sugawara T, Hisatsune C, Miyamoto H, Ogawa N & Mikoshiba K (2017). Regulation of spinogenesis in mature Purkinje cells via mGluR/PKC-mediated phosphorylation of CaMKIIβ. Proc Natl Acad Sci USA; DOI: 10.1073/pnas.1617270114.

Szegedi V, Paizs M, Csakvari E, Molnar G, Barzo P, Tamas G & Lamsa K (2016). Plasticity in Single Axon Glutamatergic Connection to GABAergic Interneurons Regulates Complex Events in the Human Neocortex ed. Bacci A. PLoS Biol 14, e2000237.

Thapliyal S, Arendt KL, Lau AG & Chen L (2022). Retinoic acid-gated BDNF synthesis in neuronal dendrites drives presynaptic homeostatic plasticity. eLife; DOI: 10.7554/elife.79863.

Topolnik L, Azzi M, Morin F, Kougioumoutzakis A & Lacaille J (2006). mGluR1/5 subtype-specific calcium signalling and induction of long-term potentiation in rat hippocampal oriens/alveus interneurones. The Journal of Physiology 575, 115–131.

Topolnik L, Chamberland S, Pelletier J-G, Ran I & Lacaille J-C (2009). Activity-Dependent Compartmentalized Regulation of Dendritic Ca^2+^ Signaling in Hippocampal Interneurons. J Neurosci 29, 4658–4663.

Topolnik L, Congar P & Lacaille J-C (2005). Differential Regulation of Metabotropic Glutamate Receptor- and AMPA Receptor-Mediated Dendritic Ca^2+^ Signals by Presynaptic and Postsynaptic Activity in Hippocampal Interneurons. J Neurosci 25, 990–1001.

Tricoire L, Pelkey KA, Daw MI, Sousa VH, Miyoshi G, Jeffries B, Cauli B, Fishell G & McBain CJ (2010). Common Origins of Hippocampal Ivy and Nitric Oxide Synthase Expressing Neurogliaform Cells. J Neurosci 30, 2165–2176.

Urban-Ciecko J, Jouhanneau J-S, Myal SE, Poulet JFA & Barth AL (2018). Precisely Timed Nicotinic Activation Drives SST Inhibition in Neocortical Circuits. Neuron 97, 611–625.e5.

Vargas-Caballero M & Robinson HPC (2004). Fast and Slow Voltage-Dependent Dynamics of Magnesium Block in the NMDA Receptor: The Asymmetric Trapping Block Model. J Neurosci 24, 6171–6180.

Vasuta C, Artinian J, Laplante I, Hébert-Seropian S, Elayoubi K & Lacaille J-C (2015). Metaplastic Regulation of CA1 Schaffer Collateral Pathway Plasticity by Hebbian MGluR1a-Mediated Plasticity at Excitatory Synapses onto Somatostatin-Expressing Interneurons. eneuro 2, ENEURO.0051-15.2015.

Walker F, Möck M, Feyerabend M, Guy J, Wagener RJ, Schubert D, Staiger JF & Witte M (2016). Parvalbumin- and vasoactive intestinal polypeptide-expressing neocortical interneurons impose differential inhibition on Martinotti cells. Nat Commun; DOI: 10.1038/ncomms13664.

Wang L & Maffei A (2014). Inhibitory Plasticity Dictates the Sign of Plasticity at Excitatory Synapses. J Neurosci 34, 1083–1093.

Wang W, Trieu BH, Palmer LC, Jia Y, Pham DT, Jung K-M, Karsten CA, Merrill CB, Mackie K, Gall CM, Piomelli D & Lynch G (2016). A Primary Cortical Input to Hippocampus Expresses a Pathway-Specific and Endocannabinoid-Dependent Form of Long-Term Potentiation. eneuro 3, ENEURO.0160-16.2016.

Weisskopf MG, Bauer EP & LeDoux JE (1999). L-type voltage-gated calcium channels mediate NMDA-independent associative long-term potentiation at thalamic input synapses to the amygdala. J Neurosci 19, 10512–10519.

Williams LE & Holtmaat A (2019). Higher-Order Thalamocortical Inputs Gate Synaptic Long-Term Potentiation via Disinhibition. Neuron 101, 91–102.e4.

Winters ND, Kondev V, Loomba N, Delpire E, Grueter BA & Patel S (2023). Opposing retrograde and astrocyte-dependent endocannabinoid signaling mechanisms regulate lateral habenula synaptic transmission. Cell Reports 42, 112159.

Yang Y & Calakos N (2013). Presynaptic long-term plasticity. Front Synaptic Neurosci; DOI: 10.3389/fnsyn.2013.00008.

